# Exploring the natural origins of SARS-CoV-2 in the light of recombination

**DOI:** 10.1101/2021.01.22.427830

**Authors:** Spyros Lytras, Joseph Hughes, Darren Martin, Arné de Klerk, Rentia Lourens, Sergei L Kosakovsky Pond, Wei Xia, Xiaowei Jiang, David L Robertson

**Affiliations:** MRC-University of Glasgow Centre for Virus Research, Glasgow, UK; Computational Biology Division, Department of Integrative Biomedical Sciences, University of Cape Town, Cape Town, South Africa; Division of Neurosurgery, Department of Surgery, Neuroscience institute, University of Cape Town, Cape Town, South Africa; Institute for Genomics and Evolutionary Medicine, Department of Biology, Temple University, Pennsylvania, USA; National School of Agricultural Institution and Development, South China Agricultural University, Guangzhou, China; Department of Biological Sciences, Xi’an Jiaotong-Liverpool University (XJTLU), Suzhou, China

**Keywords:** SARS-CoV-2, *Sarbecoviruses*, bats, origins, host range, coronaviruses, recombination, China, Southeast Asia, *Rhinolophus*, pangolins

## Abstract

The lack of an identifiable intermediate host species for the proximal animal ancestor of SARS-CoV-2, and the large geographical distance between Wuhan and where the closest evolutionary related coronaviruses circulating in horseshoe bats (*Sarbecoviruses*) have been identified, is fuelling speculation on the natural origins of SARS-CoV-2. We have comprehensively analysed phylogenetic relations between SARS-CoV-2, and the related bat and pangolin *Sarbecoviruses* sampled so far. Determining the likely recombination events reveals a highly reticulate evolutionary history within this group of coronaviruses. Clustering of the inferred recombination events is non-random with evidence that Spike, the main target for humoral immunity, is beside a recombination hotspot likely driving antigenic shift in the ancestry of bat *Sarbecoviruses*. Coupled with the geographic ranges of their hosts and the sampling locations, across southern China, and into Southeast Asia, we confirm horseshoe bats, *Rhinolophus*, are the likely SARS-CoV-2 progenitor reservoir species. By tracing the recombinant sequence patterns, we conclude that there has been relatively recent geographic movement and co-circulation of these viruses’ ancestors, extending across their bat host ranges in China and Southeast Asia over the last 100 years or so. We confirm that a direct proximal ancestor to SARS-CoV-2 is yet to be sampled, since the closest relative shared a common ancestor with SARS-CoV-2 approximately 40 years ago. Our analysis highlights the need for more wildlife sampling to (i) pinpoint the exact origins of SARS-CoV-2’s animal progenitor, and (ii) survey the extent of the diversity in the related *Sarbecoviruses*’ phylogeny that present high risk for future spillover.

**Highlights:**

- The origin of SARS-CoV-2 can be traced to horseshoe bats, genus *Rhinolophus*, with ranges in both China and Southeast Asia.
- The closest known relatives of SARS-CoV-2 exhibit frequent transmission among their *Rhinolophus* host species.
- *Sarbecoviruses* have undergone extensive recombination throughout their evolutionary history.
- Accounting for the mosaic patterns of these recombinants is important when inferring relatedness to SARS-CoV-2.
- Breakpoint patterns are consistent with recombination hotspots in the coronavirus genome, particularly upstream of the pike open reading frame with a coldspot in S1.

---

More than a year since the emergence of SARS-CoV-2, the origins of this new pandemic human coronavirus remain uncertain. First detected in association with an unusual respiratory disease outbreak in December 2019 at an animal and seafood market in Wuhan city, Hubei province, China (Li et al. 2020) no definitive progenitor of animal origin has been identified. Environmental samples taken from this market have only revealed evidence of infections linked to humans and the finding of cases with no identifiable association to this location suggests either that the market was not the epicenter for the SARS-CoV-2 spillover event, or that multiple independent spillover events occurred^1,2^. Since the 2020 coronavirus pandemic began, both generalized metagenomic and focused sampling and sequencing efforts have uncovered a number of viruses related to SARS-CoV-2, almost all retrieved from locations in China and Southeast Asia^3–8^. Several of these *Sarbecoviruses* are recombinants necessitating careful analysis as the presence of mosaic genomes violates the assumption of there being a single evolutionary history, key to reliable phylogenetic inference from mutation patterns in molecular data.

SARS-CoV-2, responsible for COVID-19, and SARS-CoV, the causative agent of the SARS outbreak in 2002-3^9^, are both members of the *Sarbecovirus* subgenus of *Betacoronaviruses*. A group of viruses which have been primarily found in horseshoe bats (family *Rhinolophidae*). Coronaviruses are known to have a chance of recombining with one another whenever these occur together within mixed infections^10,11^. Here, we comprehensively characterise the recombinant nature of the SARS-CoV-2-like coronaviruses sampled so far, focusing specifically on the phylogenetic clade of *Sarbecoviruses* that SARS-CoV-2 emerged from; hereafter referred to as the “nCoV” clade (Figure 1A)^12^. We present evidence of recombination and a handful of hotspot locations where breakpoints are over-represented, in particular upstream of Spike with a downstream coldspot (where breakpoints are under-represented) in the S1 subunit; likely a product of antigenic selection in ancestral viruses as S1 includes the two immunodominant regions NTD and RBM^13^. By comparing the phylogenies inferred for putatively non-recombinant regions of the genome (i.e., best estimates of SARS-CoV-2 and related *Sarbecoviruses* true evolutionary history) with the viruses’ sampling locations and their host’s geographic range locations, we provide a detailed understanding of the recent evolutionary histories of SARS-CoV-2’s closest known relatives including relative divergence times.

**Figure 1.**
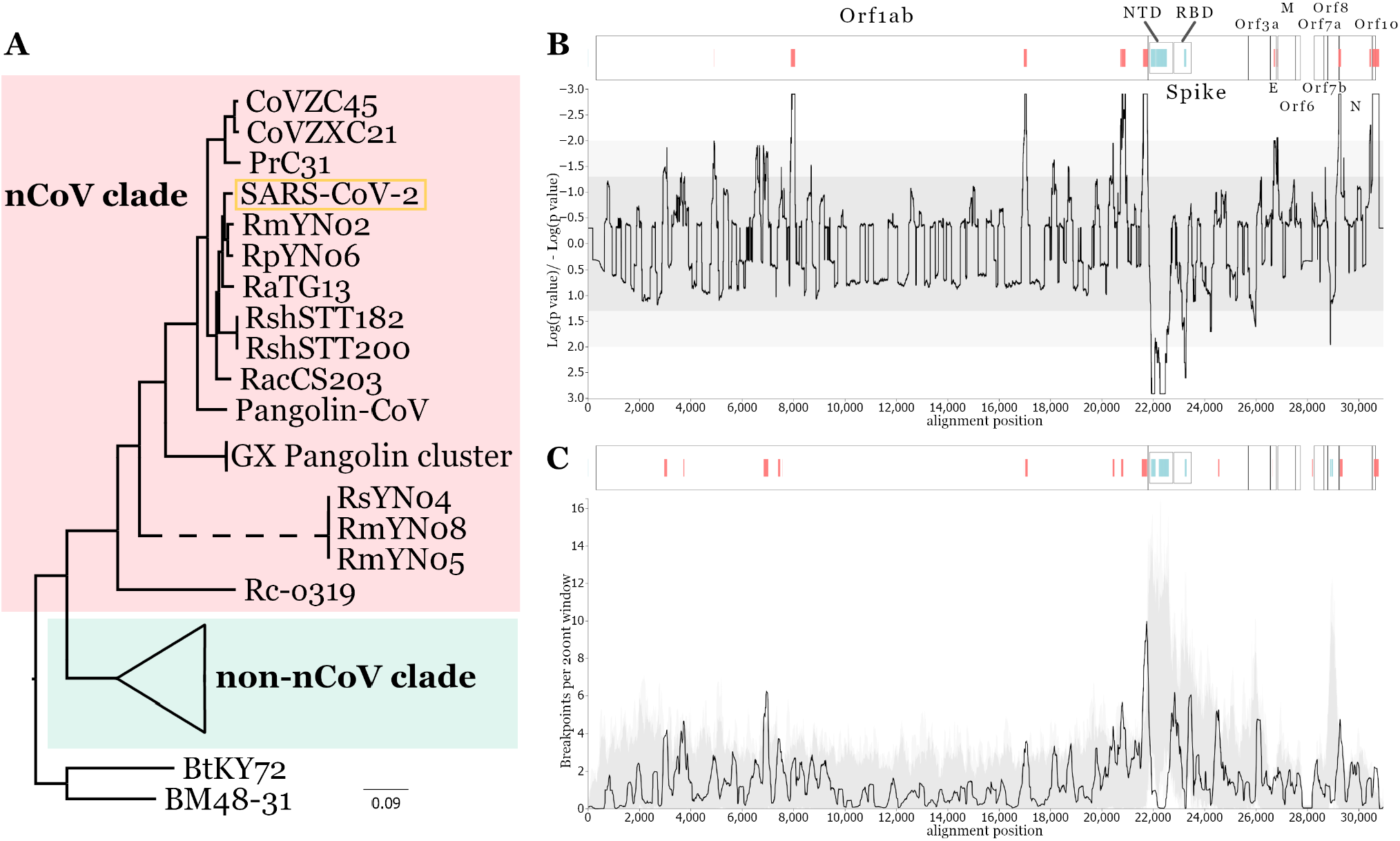
Recombination-minimised phylogeny and recombination hot-/coldspots. Maximum likelihood phylogeny inferred from a recombination-free whole genome alignment of the 78 *Sarbecoviruses* (A), see Methods. The non-nCoV/SARS-CoV clade is collapsed for clarity. All nodes presented have bootstrap confidence values above 90%. Distribution of recombination hot- and coldspots across the alignment based on the RRT (B) and the BDT (C) methods. For both plots light and dark grey represent 95% and 99% confidence intervals of expected recombination clustering. Peaks above the shaded area represent recombination hotspots and drops below represent coldspots, annotated on the corresponding ORF genome schematic above each plot by vertical red and blue lines respectively. All ORF names and the NTD and RBD encoding regions of Spike are also annotated on the schematics.

## Hotspots of recombination

For a whole-genome alignment of the set of known complete genomes from 78 *Sarbecoviruses* (including a single representative of SARS-CoV and SARS-CoV-2; Table S1) we performed an initial recombination breakpoint analysis with RDP5 (see Methods) and identified 160 unique recombination events in all the bat and pangolin-derived virus genomes. To infer a reliable phylogeny of the *Sarbecoviruses* we remove all regions with evidence for a recombination history from the genome alignment. This reconstructed non-recombinant phylogeny (Figure 1A) includes a total of 19 non-human viruses that comprise the nCoV clade that SARS-CoV-2 emerged from, a sister lineage to the non-nCoV clade SARS-CoV first emerged from in 2002.

Using the set of inferred breakpoints by RDP5, we tested for significant clustering of recombination events at specific regions of the genome, suggestive of recombination hot- or coldspots. Two permutation-based recombination breakpoint clustering tests were performed: i) a “breakpoint distribution test” (BDT) that explicitly accounts for the underlying uncertainties in the positions of identified breakpoint positions^14^ and ii) a “recombinant region test” (RRT) that focuses on point estimates of recombination breakpoint pairs that define recombination events and explicitly accounts for region-to-region variations in the detectability of recombination events^15^. Both tests provided support for the presence of several recombination hotspots: seven in the BDT and nine in the RRT analysis, assuming close locations are giving rise to the same peak (Figure 1B,C), and recombination cold-regions in the NTD and RBD domains of the Spike gene and within ORF8 (Figure 1C).

The distribution of recombination breakpoints is clearly not uniform across the *Sarbecovirus* genomes. Interestingly the pattern of hotspots near the Spike ORF has also been noted in previous research^16^. Coupled with the molecular mechanisms responsible for recombination influencing the locations of breakpoint hotspots^17^, we propose that antigenic selection and/or selection associated with switches in host receptor specificity and efficiency, i.e., antigenic shift, are the most likely candidate drivers of the observed recombination patterns. What is clear is it is imperative to account for these complex recombination patterns when examining the evolutionary history of these pathogens, since multiple evolutionary histories can be inferred from the single whole-genome alignment. As SARS-CoV-2 continues circulating in humans and mutation increases its sequence diversity, identifying SARS-CoV-2 recombination events will become easier and important to monitor^18^.

## Recombination patterns between SARS-CoV-2 relatives

To reconstruct a reliable phylogeny for a set of viruses, sufficient information needs to be present in the underlying sequence alignment. Thus, even though a whole-genome alignment can be split into shorter sub-alignments with the aim of getting rid of all independent recombination events, it is unlikely that all sub-alignments can produce reliable phylogenies. To overcome this trade-off we performed a secondary, more conservative, recombination analysis using GARD (see Methods) and identify the locations of 21 recombination breakpoints that strongly impact the inferred phylogenetic relationships of the analysed sequences when mosaic patterns are ignored. We then determined the phylogenetic relationships of the viral sequences in each of the 22 putatively non-recombinant genome regions bounded by each identifiable breakpoint (Figure 3A). The 20 nCoV viruses identified in the non-recombinant whole-genome phylogeny above (Figure 1A) were used to inform the clade annotation for the 22 new non-recombinant phylogenies.

The two genetically closest relatives of SARS-CoV-2 that were identified shortly after its emergence were the bat *Sarbecovirus*es, RaTG13 and subsequently RmYN02, both from samples collected in Yunnan^3,4^. We find RmYN02 shares a most recent common ancestor with SARS-CoV-2 about 40 years ago and RaTG13 about 50 years ago (Figure 4A) consistent with previous estimates^11,12,19^. Although SARS-CoV-2 is most similar to RmYN02 across most of its genome, the region corresponding to the first half of the RmYN02 Spike ORF appears to have been derived through recombination from a parental sequence residing outside the nCoV clade^4^ (Figure 1A). Two more viruses very recently identified in Yunnan, RpYN06^6^ and PrC31^8^ are most closely related to RmYN02 for part of their genomes. In the portion of the genome corresponding to recombination breakpoint partitioned (RBP) regions 2 to 5, the three Yunnan viruses (RmYN02, RpYN06, PrC31) cluster with strong support in a sister clade to SARS-CoV-2 (Figure 2A, Figure S1). This pattern suggests that bat sampling efforts in Yunnan have uncovered a cluster of related viruses distinct to the likely viral population of SARS-CoV-2’s proximal ancestor. Molecular dating of the RBP region 5 phylogeny (Figure 4A) indicates that this “Yunnan cluster” shared a common ancestor with SARS-CoV-2 around 1982 (95% HPD: 1970-1994). This analysis further allows us to date the node between PrC31 and RmYN02 to 2005 (95% HPD: 1998-2010), which is one of the most recent nodes in the phylogeny (Figure 4A).

**Figure 2.**
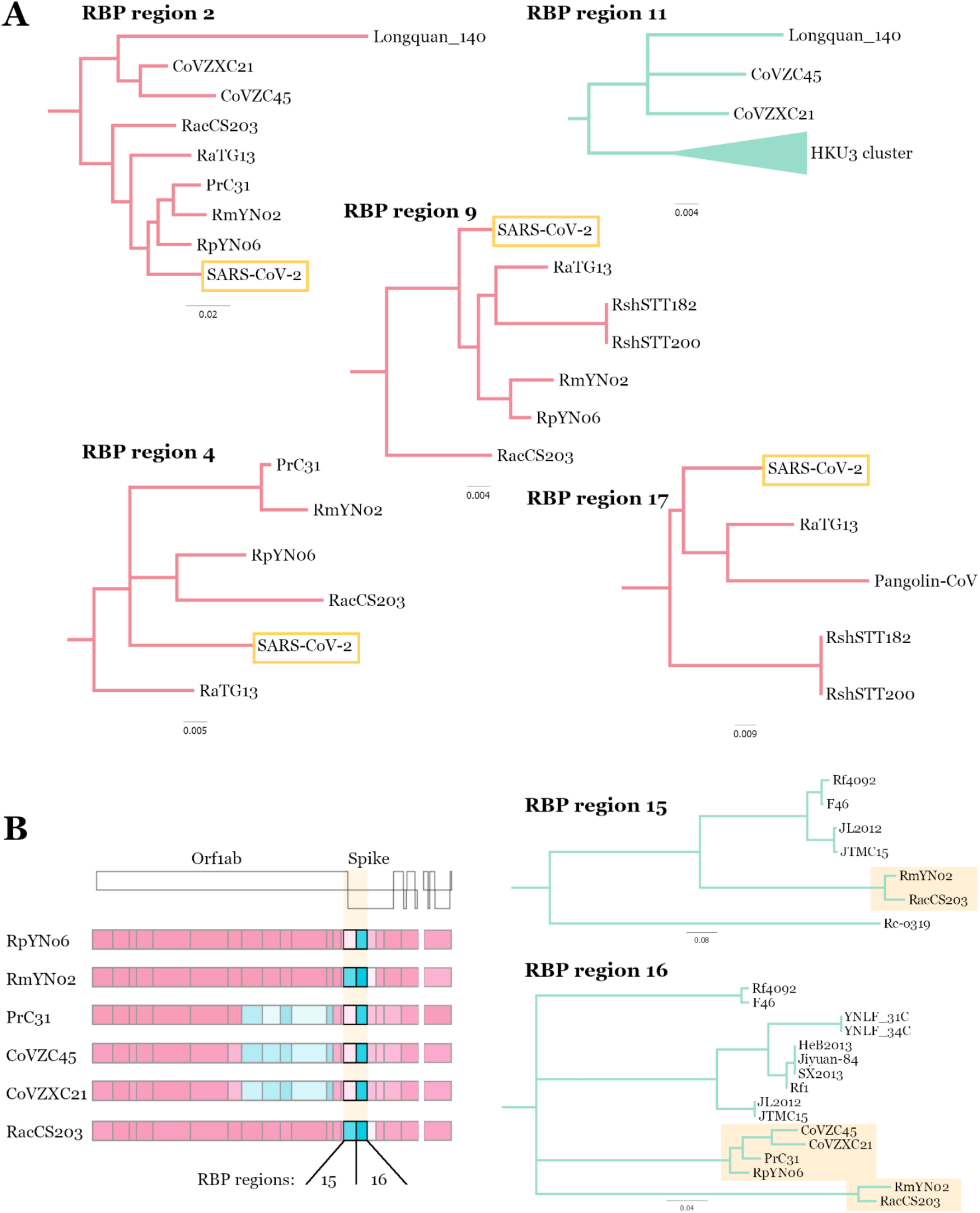
Non-recombinant topologies of SARS-CoV-2 relatives. Zoomed in regions of selected RBP region maximum likelihood phylogenies (A). Branches within the nCoV clade are coloured in red and outside the nCoV clade in green. Genome schematics of close SARS-CoV-2 relatives with recombinant Spike regions (B). RBP regions 15 and 16 are highlighted and the non-nCoV subclades of the maximum likelihood phylogenies containing the relevant viruses are presented. Nodes with bootstrap confidence values below 80% have been collapsed.

**Figure 3.**
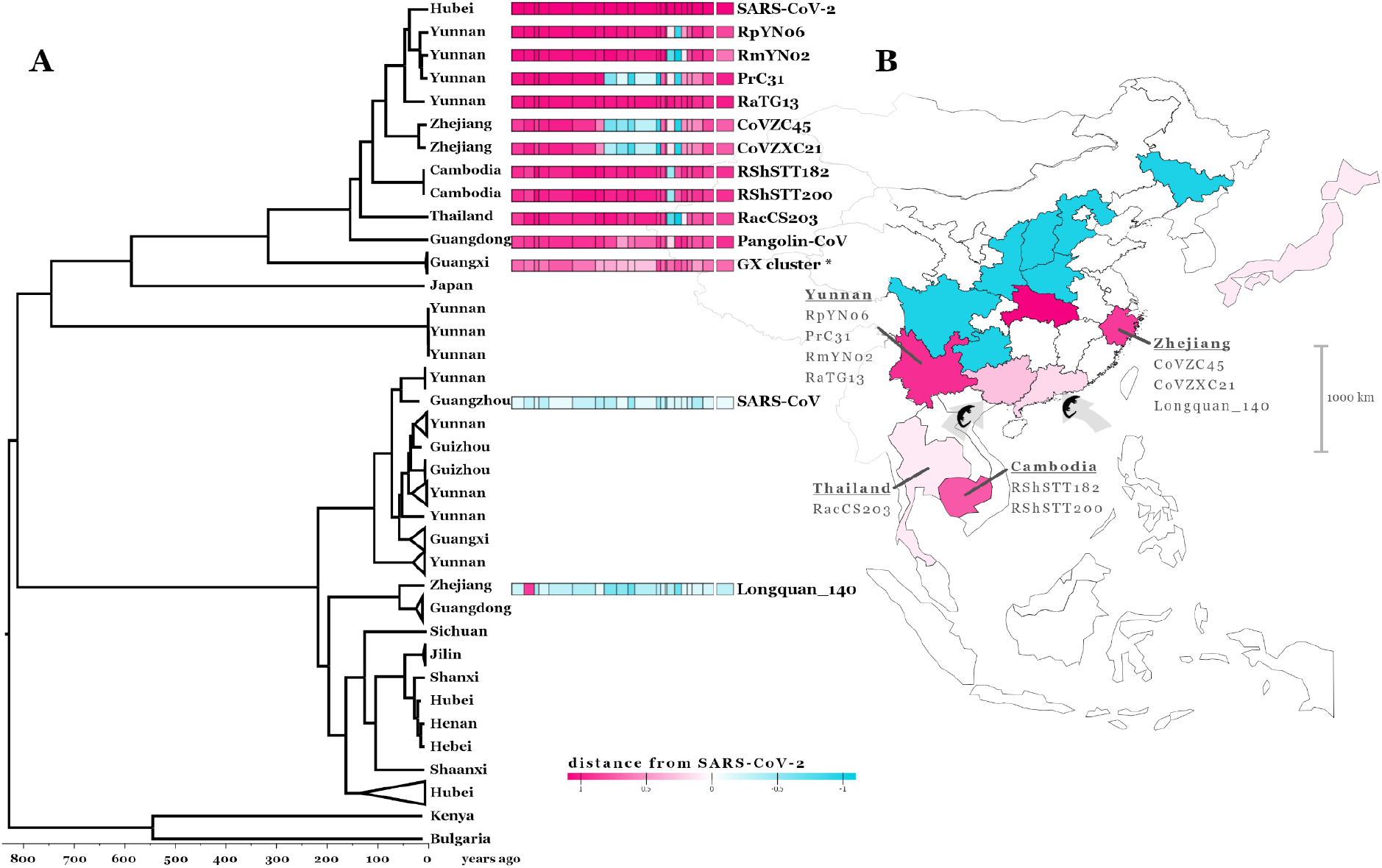
Recombination analysis and geographic distribution of *Sarbecoviruses*. Maximum clade credibility (MCC) dated phylogeny of RBP region 5 of 78 *Sarbecoviruses* (A). All tips are annotated with the geographic region the viruses have been sampled in and notable viruses are annotated with genome schematics separated into the 22 inferred RBP regions, each coloured based on phylogenetic distance from SARS-CoV-2 (see scale and Methods). RBP region 21 has been removed from the schematic due to limited phylogenetic information in the alignment. The GX cluster annotated with an asterisk contains the 5 pangolin coronaviruses collected in Guangxi. Map of East Asia with geographic regions (provinces within China, countries outside China) coloured based on *Sarbecoviruses* sampling (B): blue for regions with only non-nCoV clade samples, pink for regions where nCoV viruses have been sampled. Shading in the nCoV regions corresponds to phylogenetic distance from SARS-CoV-2 (see scale). Notable nCoV viruses and pangolin trafficking routes are annotated onto the map.

**Figure 4.**
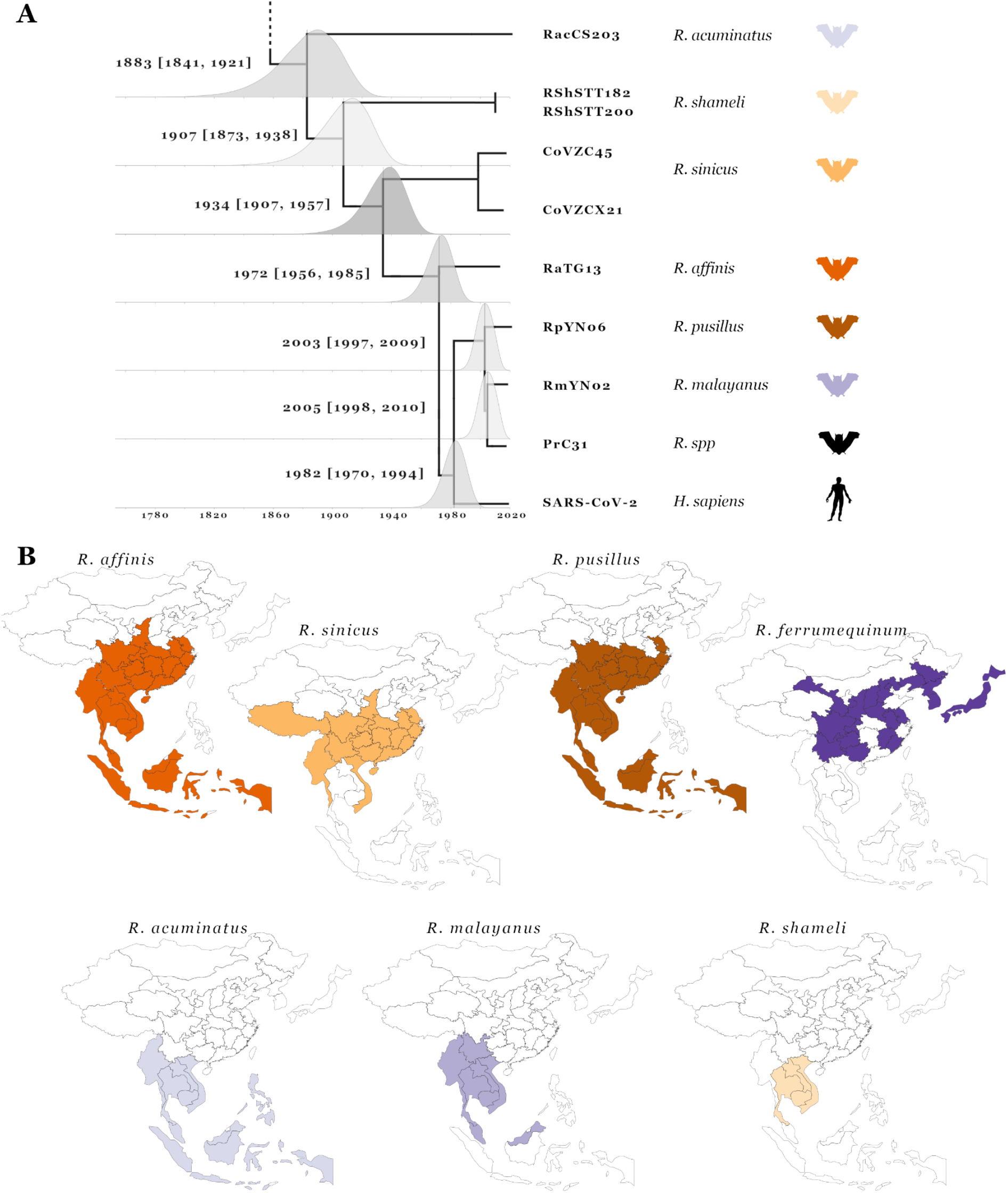
Molecular dating and *Rhinolophus* host geographic distributions. Molecular dated Bayesian phylogeny of RBP region 5 showing the 9 closest relatives to SARS-CoV-2 (A). Tree nodes have been adjusted to the mean age estimates and posterior distributions are shown for each node with mean age estimate and 95% HPD confidence intervals presented to their left. Tips are annotated with the host species they were sampled in, bat silhouette colours correspond to panel B. Geographic ranges of *Rhinolophus* species the SARS-CoV-2 closest relatives have been sampled in (B). Maps are restricted to East Asia and separated into province-level within China and country-level outside China.

The recombination analysis, however, reveals a much more complex evolutionary history for the rest of the PrC31 genome. As seen in the consensus whole-genome phylogeny (Figure 1A), most of its genome clusters with viruses CoVZC45 and CoVZXC21 sampled in Zhejiang, a coastal province in East China^20,21^. Across the majority of their genomes (excluding segments of Orf1ab and Spike) these viruses are members of the nCoV clade and share a common ancestor with SARS-CoV-2 that existed before 1934 (95% HPD: 1907-1957) according to molecular dating of RBP region 5 (Figure 4A). However, in RBP regions 8-12 the sequences of these viruses cluster outside the nCoV clade, being genetically most closely related to Zhejiang virus Longquan_140 and the HKU3 set of closely related bat *Sarbecoviruses* sampled in Hong Kong (bordering Guangdong province) (Figure 2A, Figure S1). The link between SARS-CoV-2’s closest relatives and viral populations in the southeast of South China becomes even more apparent in the phylogeny of RBP region 2 where Longquan_140 clusters within the nCoV clade along with CoVZC45 and CoVZXC21 (Figure 2A). These relationships indicate ancestral movement of the nCoV viruses across large geographic ranges in spanning Yunnan in southwest China and Zhejiang on the east coast (Figure 3B).

As more countries initiate wildlife-infecting coronavirus sampling and sequencing efforts, the geographic range of the nCoV clade linked to bat host species will be further refined, evident from the recent reporting of bat sarbecoviruses closely related to SARS-CoV-2 from: i) two samples collected in Cambodia from *R. shameli* (RShSTT182 and RShSTT200) confirmed by whole-genome analysis^7^, and ii) five bat samples from *R. acuminatus* collected in Thailand with one fully sequenced genome of virus RacCS203^5^. These viruses are, after the China sampled CoVs mentioned above, the next closest relatives to SARS-CoV-2 with common ancestor age estimates (using RBP region 5) around 1907 (95% HPD: 1873-1938) and 1883 (95% HPD: 1841-1921), respectively (Figure 4A). Similar to the other nCoV viruses, the recombination analysis uncovers more intricate phylogenetic relations for some parts of the genome. Notably, RShSTT182 and RShSTT200, despite being from Cambodia, cluster with RaTG13 for RBP regions 8 and 9 (Figure 2A, Figure S1), while in RBP region 4 of the genome RacCS203, from Thailand, clusters together with SARS-CoV-2 within the Yunnan clade (Figure 2A). This indicates that co-circulation and recombination between these viruses in the last century is responsible for the observed patterns in their inferred evolutionary history, despite the current samples having been collected across a geographic range of at least 2,500km. This wide distribution of related viruses, including shared recombination breakpoints, highlights an important feature of bat species: their frequently overlapping/sympatric ranges will provide ample opportunities for transmissions of viral variants from one bat species (or sub-species) to another.

Consistent with the Spike S1 recombination hotspots revealed in the initial analysis (Figure 1B,C), most close relatives of SARS-CoV-2 presented here have non-nCoV derived recombinant sequences at the start of the Spike gene (Figure 2B). Despite one collected from Yunnan, China and the other from Cambodia, viruses RmYN02 and RacCS203 share a closely related non-nCoV sequence in RBP regions 15 and 16 (encompassing the Spike NTD and RBD respectively; Figure 2B) having a distinct RBD compared to that of SARS-CoV-2^4,5^. On the other hand, viruses RpYN06, PrC31, CoVZC45 and CoVZXC21 cluster within the nCoV clade for region 15 but move into broadly the same non-nCoV cluster as RmYN02 and RacCS203 for region 16 (Figure 2B). We speculate that some of the apparent patterns of recombination-mediated exchange between nCoV and non-nCoV viruses can be partly explained by recombination “overprinting”. This will occur when an nCoV virus has acquired a non-nCoV genomic sequence through ancestral recombination but its progenitors co-circulating with other nCoV viruses incurred subsequent recombination events that overlapped portions of the original non-nCoV recombinant sequence, producing the more complex “patchy” patterns we see in the currently sampled viruses. Note, overprinting of recombination regions will result in reduced confidence in the breakpoints at deeper nodes in the phylogeny.

The finding that Sunda (also known as Malayan) pangolins, *Manis javanica*, non-native to China, are the other mammal species from which nCoV *Sarbecoviruses* have been sampled in Guangxi and Guangdong provinces in South China^22,23^, indicates these animals are likely being infected in this part of the country (Figure 3B). Pangolins are one of the most frequently trafficked animals with multiple smuggling routes leading to southern China^24^. The most common routes involve moving the animals from Southeast Asia (Myanmar, Malaysia, Laos, Indonesia, Vietnam) to Guangxi, Guangdong, and Yunnan. The most likely scenario that is consistent with both the respiratory distress that the sampled pangolins exhibited^23,25^ and the lack of evidence of similar infections among Sunda pangolins in Malaysia^26^, is that the viruses obtained from these animals infected them (presumably from bat sources) after they were trafficked into southern China.

Although the recombination patterns evident in the pangolin-derived virus genomes seem to be less complex than those of the bat nCoV genomes, the Guangdong Pangolin-CoV has a Spike receptor binding domain that is most similar to that of SARS-CoV-2, likely due to recombination in the other sampled close relatives, e.g., RaTG13^11^ (and reflected in region 17, Figure 2A). The susceptibility of pangolins to an apparently new human coronavirus is not surprising given the well-documented generalist nature of SARS-CoV-2^27^, which has been found to readily transmit to multiple mammals with similar ACE2 receptors and poses a grave risk of reverse-zoonoses as has been most notably demonstrated with human to mink transmissions^28^.

## Overlapping horseshoe bat ranges

Considering that almost all *Sarbecoviruses* have been sampled in related bat hosts, i.e., the *Rhinolophidae*, horseshoe bats, which have ranges that span different regions where nCoV clade viruses have been collected (Figure 4B), should be prioritized for sampling. For example, the intermediate horseshoe bat species, *R. affinis*, is sufficiently dispersed across China to account for the geographical spread of i) bat *Sarbecovirus* recombinants in the West and East of China, ii) infected imported pangolins in the South, iii) bat *Sarbecovirus* recombinant links to southwest of China, and iv) SARS-CoV-2 emergence towards Hubei in Central China (Figure 3B). Strikingly, the ranges of multiple species including *R. affinis, R. sinicus* and *R. pusillus* overlap all the regions of China where the nCoVs have been collected (Figure 4B). Other horseshoe bat species that might harbour nCoVs have different ranges, for example, *R. ferrumequinum*, being absent from large parts of South Central China, and *R. malayanus* found predominantly in the western part of China and countries to the Southwest of China (Myanmar, Thailand, Cambodia, Laos, Viet Nam, and Peninsular Malaysia)^29^. RmYN02, RmYN05 and RmYN08 were sampled from *R. malayanus* in Yunnan^4,6^ (Figure 4B) indicating that the nCoV viruses are being exchanged between bat species in regions where ranges overlap such as Yunnan, linking hosts found predominantly in China with hosts in the Indochinese peninsula.

The wide geographic ranges of *R. pusillus* and *R. affinis* and the fact that two of the closest known relatives of SARS-CoV-2, RpYN06 and RaTG13, have been sampled in these species flags them as prime suspects for the source of the SARS-CoV-2’s progenitor in China. Additionally, these two bat species are found in shared roosts with *R. sinicus* and *R. ferrumequinum* in Yunnan and with *R. sinicus* in Guangxi^30^, providing opportunities for host switches, co-infections and thus recombination between the sarbecoviruses that these bat species carry. *R. pusillus* and *R. affinis* also link more regions of China with bat species such as *R. shameli, R. malayanus* and *R. acuminatus* which are only found in Southeast Asia and southwest of China (Figure 4B). Latinne et al. (2020)^31^ recently published a large-scale sampling expedition of coronaviruses across bats in China. Although only short RdRp fragments were sequenced, reconstructing the phylogeny for the novel viruses reveals a cluster of seven identical *Sarbecovirus* sequences sampled from *R. affinis* within the nCoV clade, that cluster close to SARS-CoV-2 (Figure S2). Still, the fact that viruses in the Yunnan clade (consisting of RmYN02, RpYN06 and PrC31) were sampled from three different *Rhinolophus* species supports the hypothesis that these viruses readily infect multiple different horseshoe bat species with overlapping geographical ranges.

Based on the analysis of the *Sarbecovirus* and host data presented here, we propose that horseshoe bat population sampling should focus on the known ranges of probable bat hosts. Specifically, samples should be collected in roosting environments spread across China with care being taken both to avoid a further spillover or reverse zoonosis and to protect the bat species^30^. Sampling strategies will also need to consider the distinct subspecies of *Rhinolophus* as the delineators of genetically meaningful host populations for coronaviruses. For example, there are two *R. affinis* sub-species on mainland China: *himalayanus* and *macrurus*^32^. Future sampling should also encompass a range of indigenous mammals other than bats that we now know can be infected by coronaviruses. Although highly endangered, it is possible that Chinese pangolins, given their susceptibility to infection and their geographical range across southern China^33^, might be the “missing” intermediate host of the SARS-CoV-2 proximal ancestor^1^.

## Conclusion

The currently available data, although sparse, illustrates a complex reticulate history involving the lineage of *Sarbecoviruses* SARS-CoV-2 emerged from, governed by co-circulation of related coronaviruses, over at least the last 100 years, across the bat populations in southern China, and into Southeast Asia with multiple recombination events imprinted on the genomes of these viruses. Considering the high frequency of recombination, it is expected that selection could preferentially favour exchanges of specific genome regions, in line with our detection of hotspots near the Spike gene (Figure 1B,C). Our analysis further illustrates the importance of accounting for recombination rather than using whole-genome pairwise similarity to determine the shared evolutionary history of these viruses. This is exemplified by RaTG13 which is often described as the “on average” closest *Sarbecovirus* to SARS-CoV-2 despite not being the closest virus once recombination history is accounted for in the other nCoV *Sarbecoviruses* (Figure 1A, 3A).

The evidence of recombination events between *Sarbecoviruses* sampled in different geographical regions and from different bat hosts, indicates recent extensive movement of the viruses between different regions and species (and presumably sub-species too) as a result of the different bat populations that carry them coming into contact with one another. Although very few nCoV viruses are known to be hosted by mammals other than horseshoe bats, the recombination patterns detected within the nCoV genomes imply the existence of one or a few primary reservoir hosts with a geographical range spanning Thailand from the Southwest and Zhejiang to the East, a distribution that is consistent with specific Chinese horseshoe bats being the primary reservoir hosts. Having presented evidence in support of *R. affinis* and *R. pusillus*’s potential significance as the reservoir species, it should be noted that at least 20 different *Rhinolophus* species are distributed across China (four being endemic to China), many of which have not yet been found hosting nCoVs. The generalist nature of *Sarbecoviruses* also means multiple wild or farmed animals (e.g., minks)^**1**^ could have facilitated transmission of SARS-CoV-2 from bats to humans.

The risk of future emergence of a new SARS-CoV-2 like nCoV strain in humans is too high to restrict sampling strategies. Beyond the relatively rare detection of SARS-like antibodies in people from rural communities in China^34,35^, SARS related coronaviruses have not, to our knowledge, seeded outbreaks in humans before. This indicates that there is limited human exposure to these viruses, suggesting ecological “barriers” to their emergence^36^. One possible recent disruption that caused widespread and unusual movements of animals in China and may have increased the permeability of this barrier, was the dramatic shortage of pork products in 2019^37^ attributable to an African swine fever virus (ASFV) outbreak impacting 100s of millions of pigs in China^38^. Such a major disruption in the food supply chain will have potentially brought humans into increased contact with *Sarbecovirus* infected animals as i) exotic meats replaced pork, ii) animals from rural locations were brought to city markets and/or iii) by meat from infected animals being transported in cold chain processes. Further facilitating animal contacts, increased human encroachment into rural areas as a result of new and faster travel networks around and between metropolitan areas have also likely increased opportunities for *Sarbecoviruses* to spillover into humans. Given the reality of frequent human-animal contact, routine characterisation of respiratory infections would seem a sensible precaution to prevent future emergence of *Sarbecoviruses*.

The key and most urgent questions relating to the prevention of another emergence, is thus **not** how did SARS-CoV-2 get from Yunnan to Hubei, but rather which bat or other animal species are harbouring nCoV *Sarbecoviruses* and what are the risks of a future spillover? There is undoubtedly a virus highly related to SARS-CoV-2 still present somewhere most probably in a bat species in South China or towards Yunnan in the southwest. The best we can do is maximize the probability that future sampling efforts will uncover that host species or sub-species.

## Methods

### Genome alignment

The whole genome sequences of the 78 *Sarbecoviruses* used in this analysis (Table S1) were aligned and the open reading frames (ORF) of the major protein-coding genes were defined based on SARS-CoV-2 annotation^39^. To minimise alignment error codon-level alignments of the ORFs were created using MAFFT v7.453^40^ and PAL2NAL^41^. The intergenic regions were also aligned separately using MAFFT and all alignments were pieced together into the final whole-genome alignment and visually inspected in Bioedit^42^.

### Genome-specific recombination analysis

We first performed an analysis for detecting individual recombination events in individual genome sequences using the RDP^43^, GENECONV^44^, BOOTSCAN^45^, MAXCHI^46^, CHIMAERA^47^, SISCAN^48^, and 3SEQ^49^ methods implemented in the program RDP5^50^. Default settings were used throughout except: i) only potential recombination events detected by three or more of the above methods, coupled with phylogenetic evidence of recombination were considered significant and ii) sequences were treated as linear. Using the RDP5 approach l (http://web.cbio.uct.ac.za/~darren/rdp.html), the approximate breakpoint positions and recombinant sequence(s) inferred for every potential recombination event, were manually checked and adjusted where necessary using the phylogenetic and recombination signal analysis features available in RDP5. Breakpoint positions were classified as undetermined if the 95% confidence interval on their location overlapped: i) the 5′ and 3′ ends of the alignment; or ii) the position of another detected breakpoint within the same sequence (in such cases it could not be discounted that the actual breakpoint might not have simply been lost due to a more recent recombination event). All of the remaining breakpoint positions were manually checked and adjusted when necessary using the BURT method with the MAXCHI matrix and LARD two breakpoint scan methods^51^ used to resolve ties. A putatively non-recombinant version or the original whole-genome alignment was reconstructed by excluding all minor parent sequence segments based on the RDP5 analysis.

### Recombination hotspot analysis

The distribution of 236 unambiguously detected breakpoint positions defining 160 unique recombination events based on the RDP5 analysis described above were analysed for evidence of recombination hot- and cold-spots using the permutation-based “recombinant region test” (RRT)^15^ and “breakpoint distribution test” (BDT)^14^. The RRT accounts for site-to-site variations in the detectability of individual recombination events and examines the distribution of point estimates of pairs of breakpoint locations bounding each of the unique recombination events detected by RDP5. Rather than using point estimates of recombination breakpoint locations, the BDT accounts for underlying uncertainties in the estimation of individual breakpoint locations as determined from the state transition likelihoods yielded by the hidden Markov model-based recombination breakpoint detection method, BURT (described in the RDP5 program manual at http://web.cbio.uct.ac.za/~darren/rdp.html).

### Whole-genome alignment recombination analysis

Next, we sought to conservatively examine the entire genome alignment for recombination breakpoints using the Genetic Algorithm for Recombination Detection (GARD) method^52^ implemented in Hyphy v2.5.29^53^. Likelihood was evaluated using the Akaike Inference Criterion (AIC)^54^. To improve computational efficiency and focus on the closest relatives of SARS-CoV-2 22 of the 78 viruses that are closest to SARS-CoV-2 or had preliminary evidence of clustering near between clade recombinants were included in the GARD analysis (Table S1). Only breakpoint present in more than 2/3 of the 64 GARD consecutive models were retained to produce a final set of 21 likely breakpoints (positions corresponding to the SARS-CoV-2 reference genome Wuhan-Hu-1 in order: 1680, 3093, 3649, 4973, 8208, 11445, 12622, 14401, 15954, 16923, 19965, 20518, 21198, 21411, 22460, 23396, 24144, 24843, 26323, 27388, 27685). Based on these the whole-genome alignment was split into 22 recombinant breakpoint partitioned (RBP) regions.

### Phylogenetic reconstruction

The phylogeny of each RBP alignment region based on the GARD analysis and the non-recombinant whole-genome based on the RDP5 analysis were reconstructed using iqtree version 1.6.12^55^ under a general time reversible (GTR) substitution model assuming invariable sites and a 4 category Γ distribution. Tree node confidence was determined using 10,000 ultrafast bootstrap replicates.

Based on the non-recombinant whole-genome phylogeny, 20 viruses form a monophyletic nCoV clade (Figure 1A). To illustrate the distance of each virus from SARS-CoV-2 for each GARD determined genomic region, we defined the nCoV clade on each phylogeny as the subset of the aforementioned 20 nCoV viruses forming a monophyly with SARS-CoV-2 in each phylogeny. The rest of the viruses were classified as members of the non-nCoV clade for each RBP region. We then used an arbitrary tip distance scale normalised between all phylogenies so distances are comparable between regions. For each maximum likelihood tree, the tip distance between each tip and SARS-CoV-2 is calculated using ETE 3^56^ as d1 for members of the nCoV clade and d2 for members of the non-nCoV clade. The distances are then normalised so that for nCoV clade members range between 0.1 and 1.1 (1.1 being SARS-CoV-2 itself and 0.1 being the most distant tip from SARS-CoV-2 within the nCoV clade) and between -0.1 and -1.1 for non-nCoV members (−0.1 being the closest non-nCoV virus to SARS-CoV-2 and -1.1 the most distant), as follows:

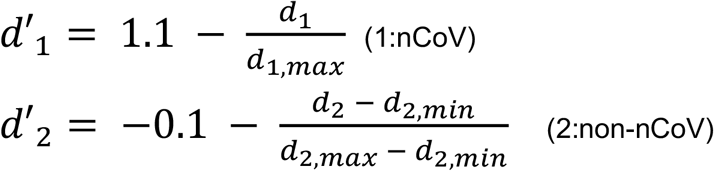

With d’1 and d’2 being the normalised values for each clade, variables denoted with “min” being the smallest distance and variables denoted with “max” being the largest distance in each given set.

Phylogenies were visualised using FigTree (http://tree.bio.ed.ac.uk/software/figtree/) and ETE 3^56^.

### Molecular dating

To provide temporal information to the phylogenetic history of the viruses, we performed a Bayesian phylogenetic analysis on RBP region 5, using BEAST v1.10.4^57^. This region was selected due to its length, being one of the two longest non-recombinant regions in the analysis (3,238 bp), and because all 20 nCoV viruses form a monophyly in the respective tree. Based on the observation of an increased evolutionary rate specific to the deepest branch of the nCoV clade reported in MacLean et al. (2020)^12^, we adopted the same approach of fitting a separate local clock model to that branch from the rest of the phylogeny. A normal rate distribution with mean 5×10^−4^ and standard deviation 2×10^−4^ was used as an informative prior on all other branches. The lineage containing the BtKY72 and BM48-31 bat viruses was constrained as the outgroup to maintain overall topology. Codon positions were partitioned and a GTR+Γ substitution model was specified independently for each partition. The maximum likelihood phylogeny reconstructed previously for RBP region 5 was used as a starting tree (rooted at the BtKY72 and BM48-31 clade). A constant size coalescent model was used for the tree prior and a lognormal prior with a mean of 6 and standard deviation of 0.5 was specified on the population size. Two independent MCMC runs were performed for 500 million states for the dataset. The two chains were inspected for convergence and combined using LogCombiner^58^ using a 10% burn-in for each chain. The effective sample size for all estimated parameters was above 200.

### Host range data

All host ranges presented in Figure 4B were retrieved from the IUCN Red List of Threatened Species (https://www.iucnredlist.org/)^29^ and the Mammals of China (Princeton Pocket Guide)^59^. Geographic visualisation was performed using D3 and JavaScript in Observable (https://observablehq.com/).

## Acknowledgements

We would like to thank all the authors who have kindly deposited and shared genome data on GISAID. Credit also needs to be given to the surveillance projects for generating the genome data that is available in GenBank and to the software developers for making the tools we have used freely available. A table with genome sequence acknowledgments can be found in supplementary material. DLR and JH are funded by the MRC (MC_UU_1201412) and DLR by the WT (220977/Z/20/Z). SL is funded by an MRC studentship. DPM is funded by the Wellcome Trust (222574/Z/21/Z).

## Supplementary

**Figure S1**. Maximum likelihood phylogenies for all 22 RBP regions of the analysis. The nCoV clade is annotated in pink and the non-nCoV clade in blue. SARS-CoV-2 and SARS-CoV are highlighted in pink and blue respectively. Branch length (top) and bootstrap support (bottom) are shown on every node. Nodes with support below 80 have been collapsed.

**Figure S2**. Maximum likelihood phylogeny reconstructed using iqtree (GTR+I+Γ4) of all 78 *Sarbecoviruses* used throughout the analysis, including the short RdRp fragments of related *Sarbecoviruses* reported in Latinne et al. (2020)^31^. The genomic region used for the alignment corresponds to the SARS-CoV-2 reference genome’s Wuhan-Hu-1 coordinates 15280 - 16282. Nodes with bootstrap support (10,000 replicates) below 80 have been collapsed. The nCoV clade is annotated in pink and the non-nCoV clade in blue. SARS-CoV-2 and SARS-CoV are highlighted in pink and blue respectively. Viruses from Latinne et al. are highlighted in grey, apart from the 7 sequences that cluster within the nCoV clade which are highlighted in green. Out of this cluster of sequences MN312634.1 has been collected from a confirmed *R. affinis* bat species.

**Table S1**. Accessions, metadata and GISAID acknowledgments for all 78 virus genomes used in this analysis.

