## Supplementary figures S1 and S2, and table S1. for "Exploring the natural origins of SARS-CoV-2 in the light of recombination": Lytras-etal_nCoV_origins-suppFigsS1-S2.pdf

Non-recombinant region 1

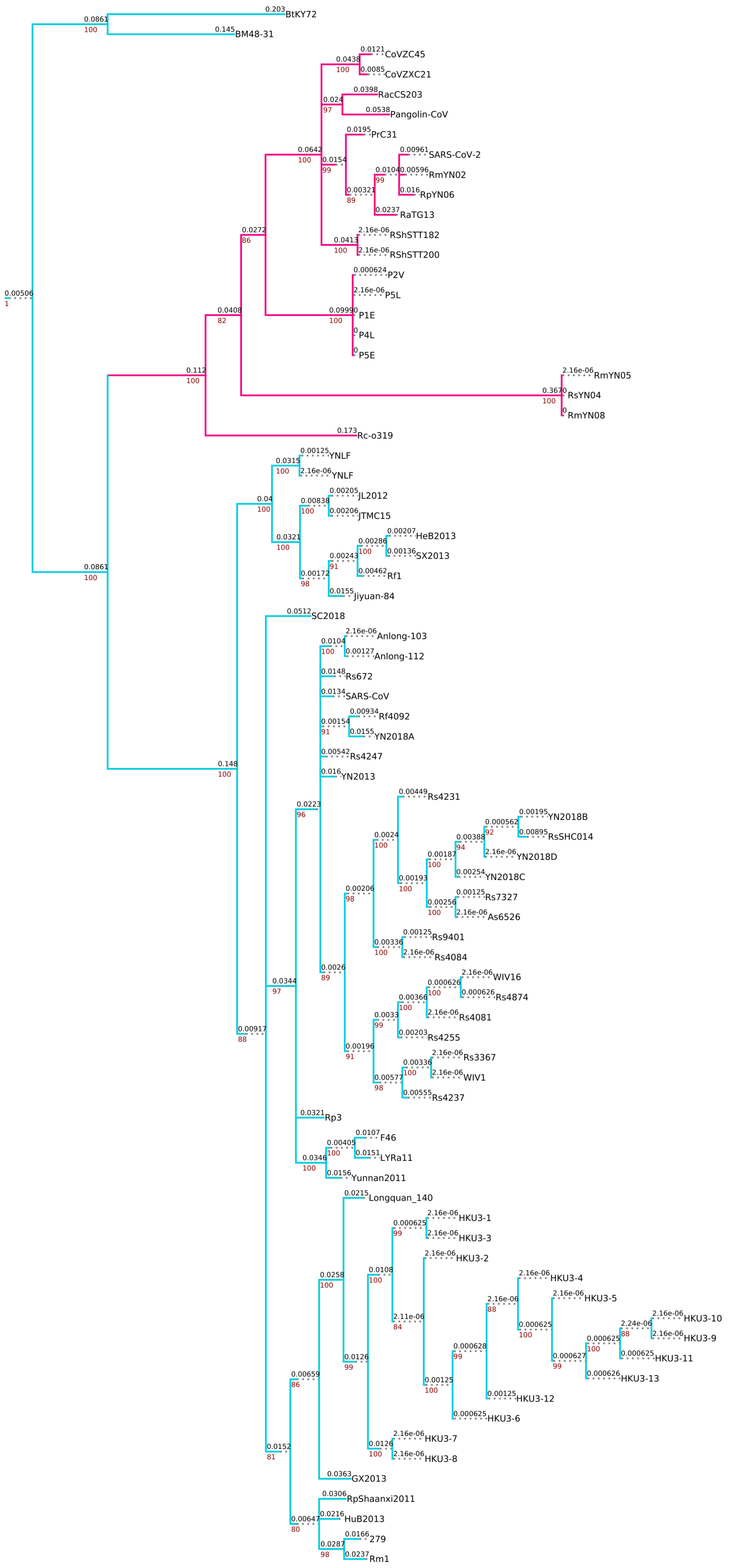


**Figure S1.** Maximum likelihood phylogenies for all 22 RBP regions of the analysis. The nCoV clade is annotated in pink and the non-nCoV clade in blue. SARS-CoV-2 and SARS-CoV are highlighted in pink and blue respectively. Branch length (top) and bootstrap support (bottom) are shown on every node. Nodes with support below 80 have been collapsed.

Non-recombinant region 2

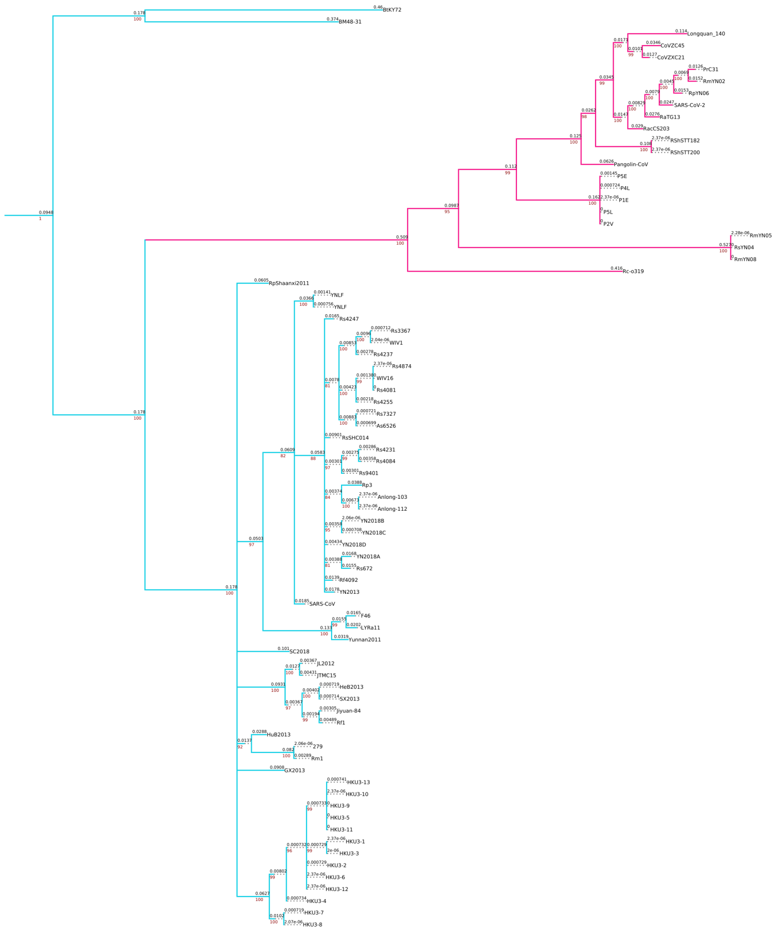


**Figure S1. continued.**

Non-recombinant region 3

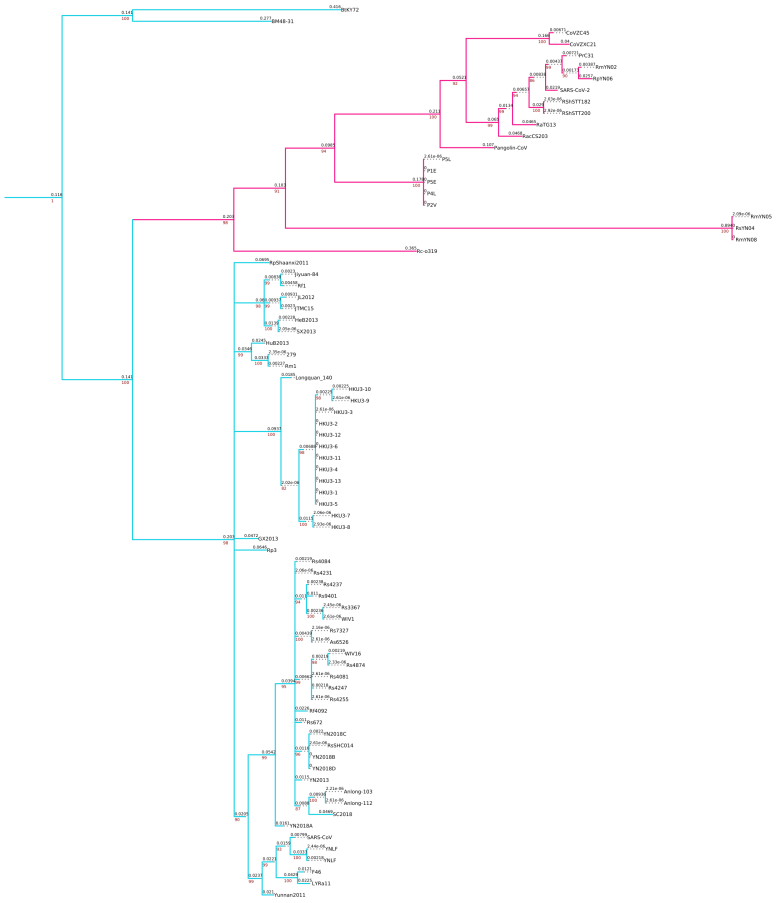


**Figure S1. continued.**

Non-recombinant region 4

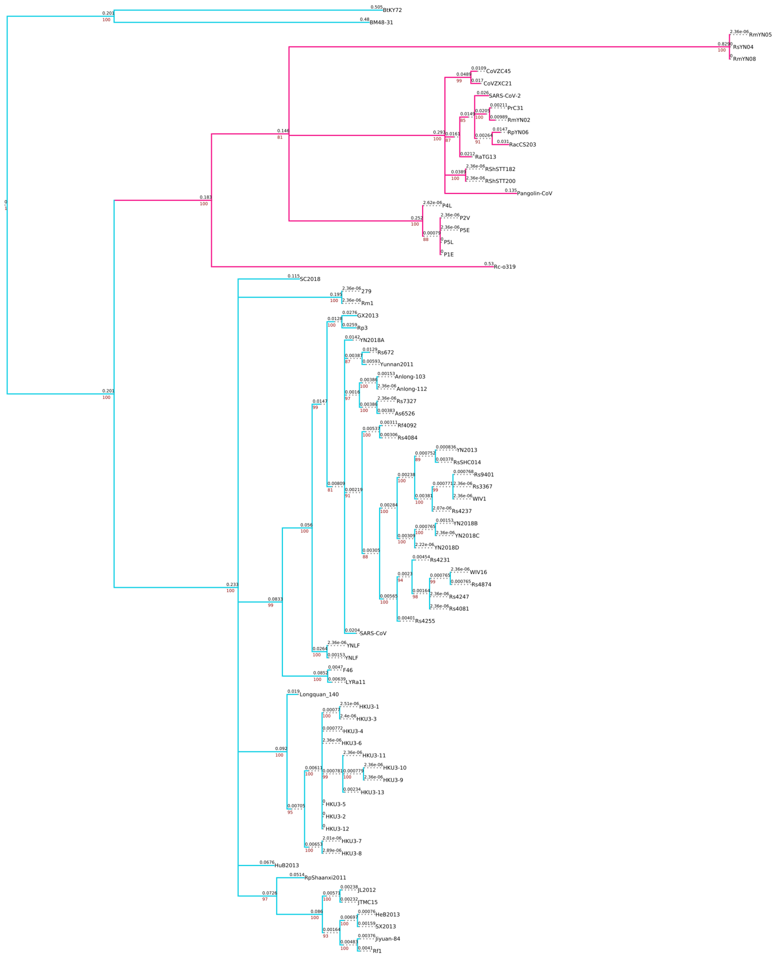

**Figure S1. continued.**

### Non-recombinant region 5

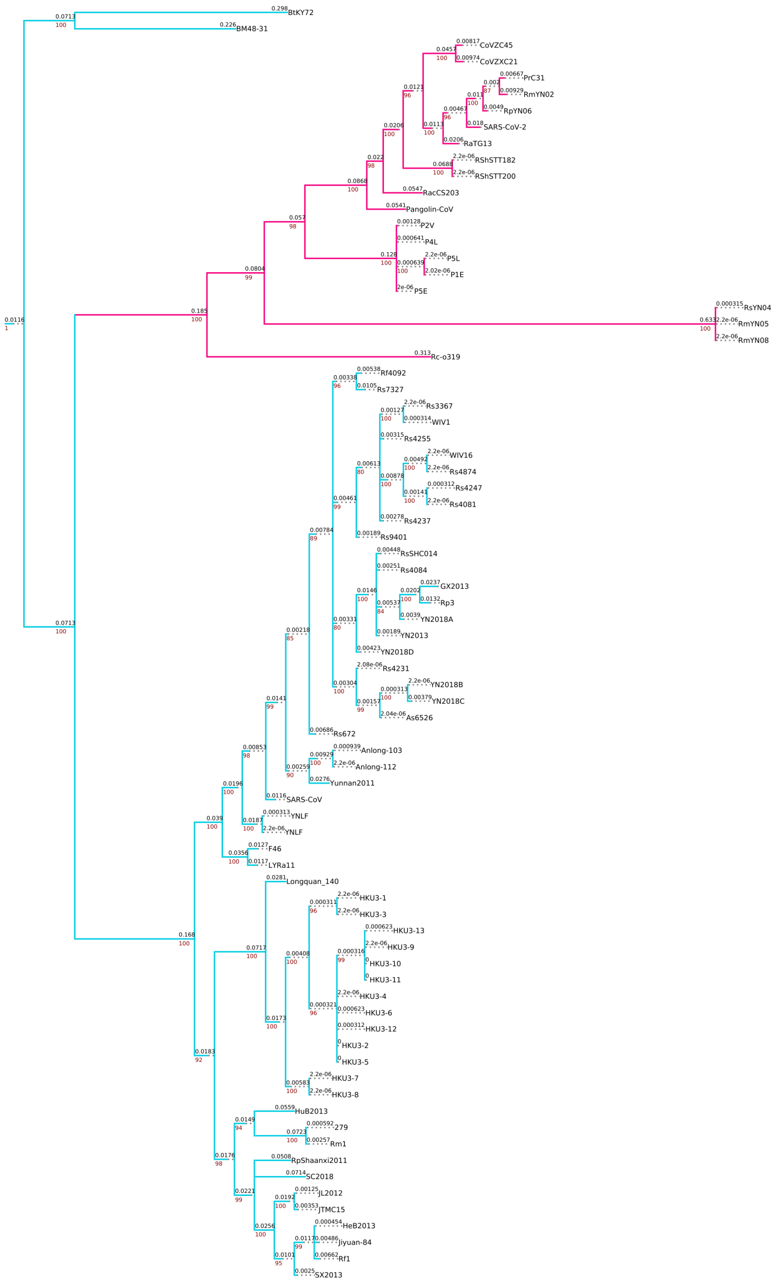


**Figure S1. continued.**

Non-recombinant region 6

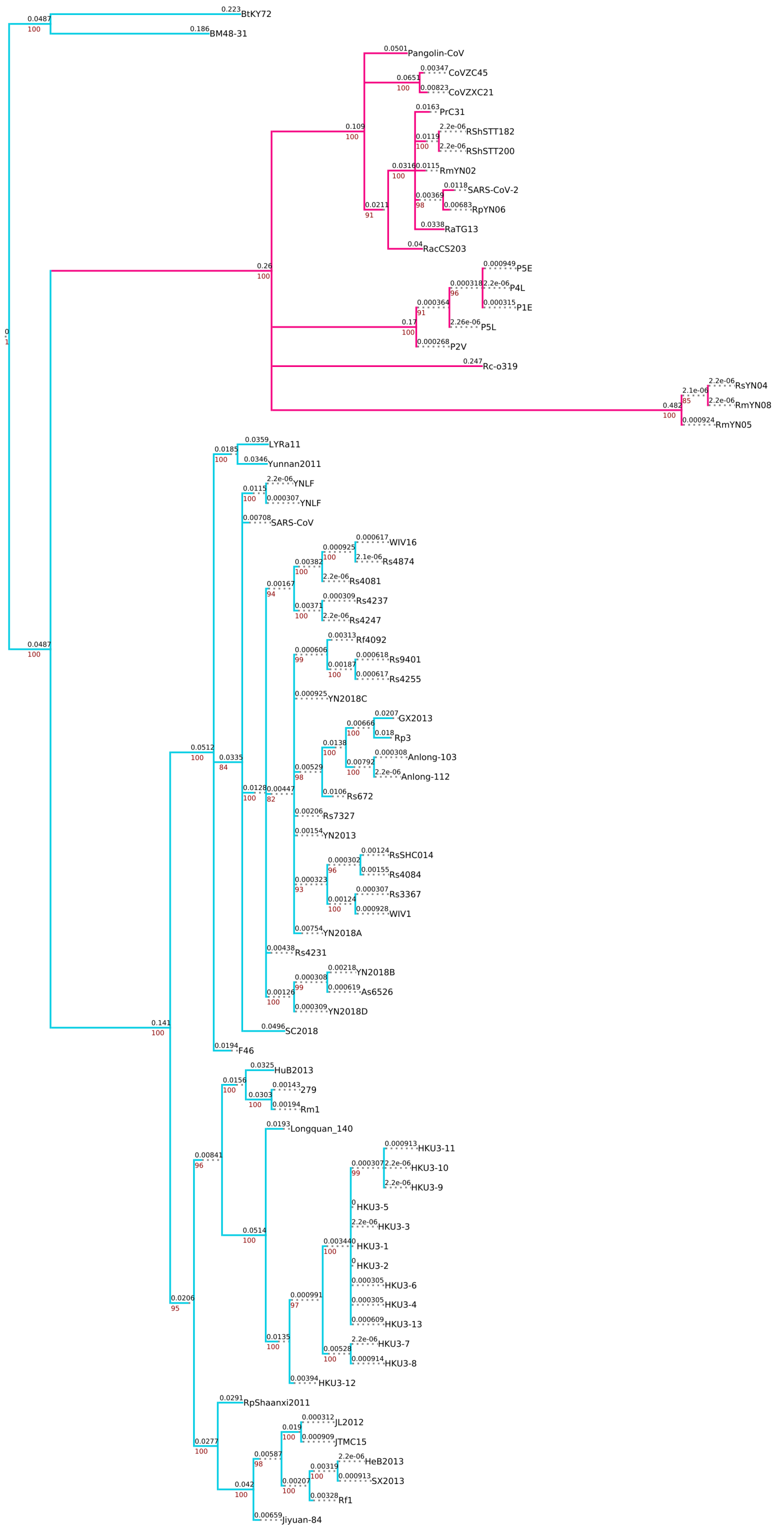


**Figure S1. continued.**

Non-recombinant region 7

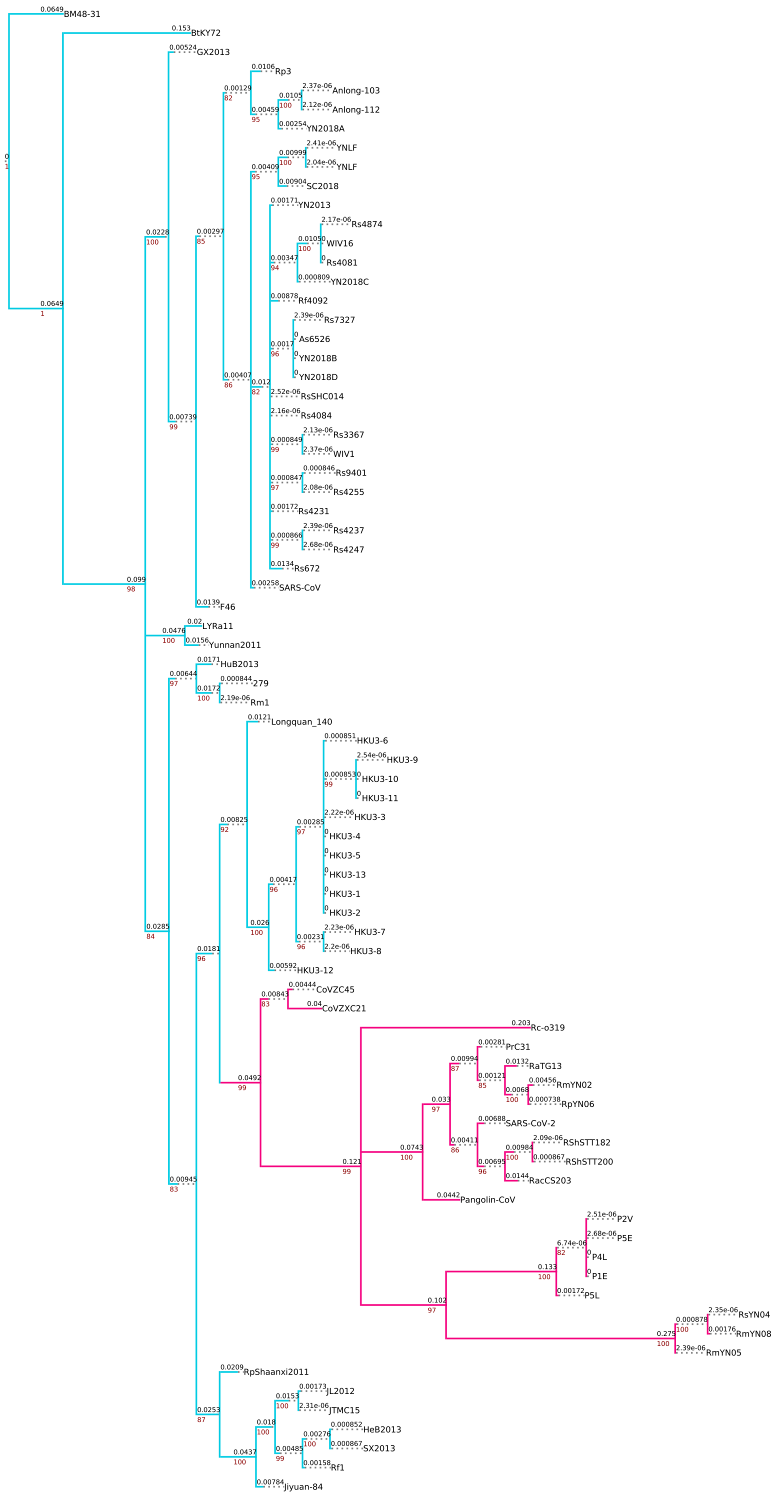

**Figure S1. continued.**

Non-recombinant region 8

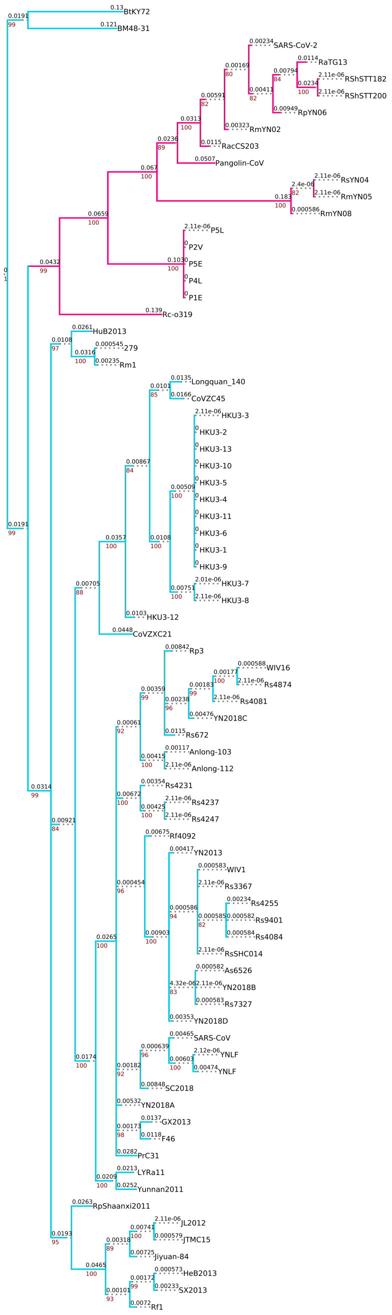


**Figure S1. continued.**

Non-recombinant region 9

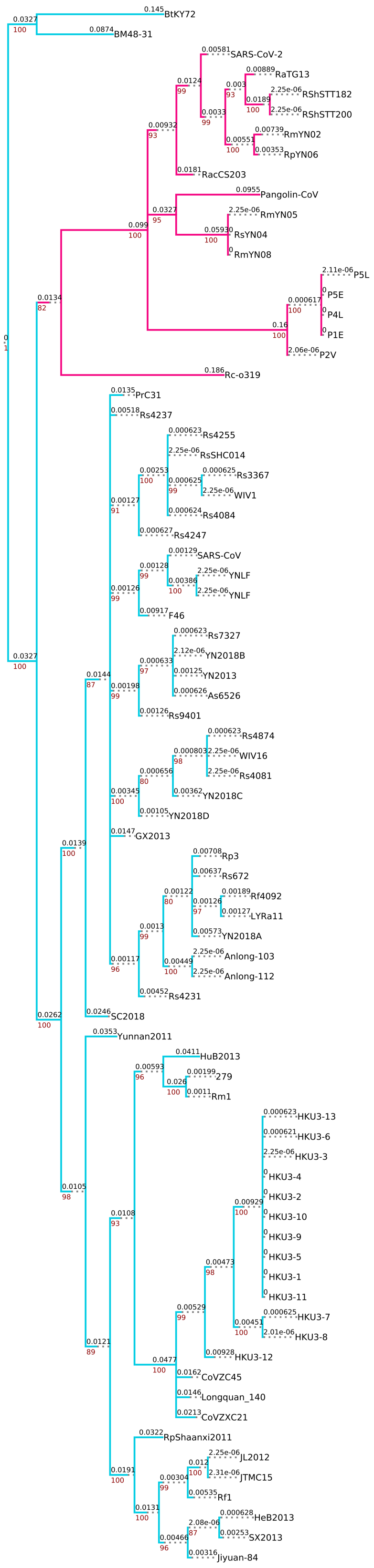


**Figure S1. continued.**

Non-recombinant region 10

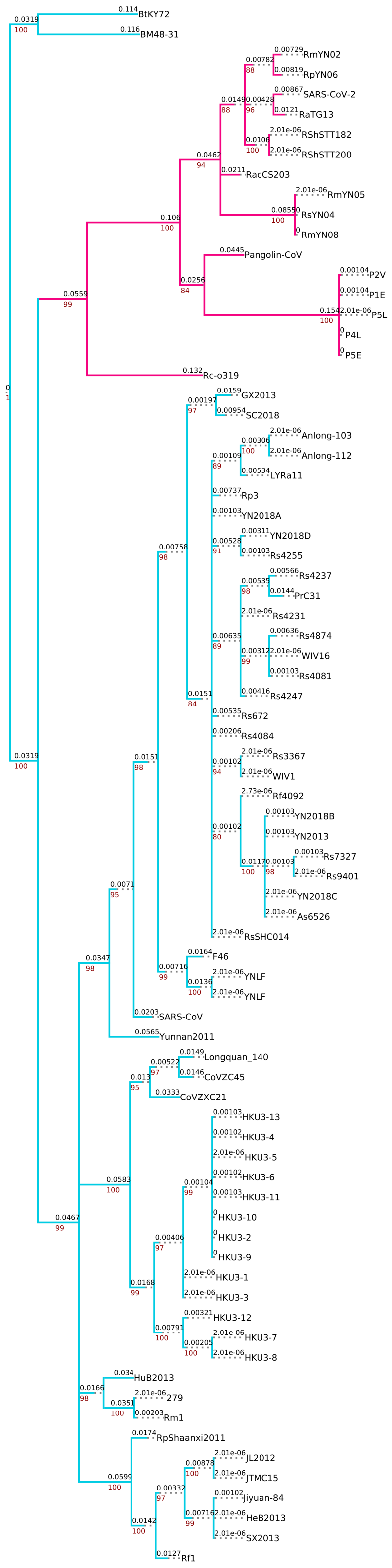


**Figure S1. continued.**

Non-recombinant region 11

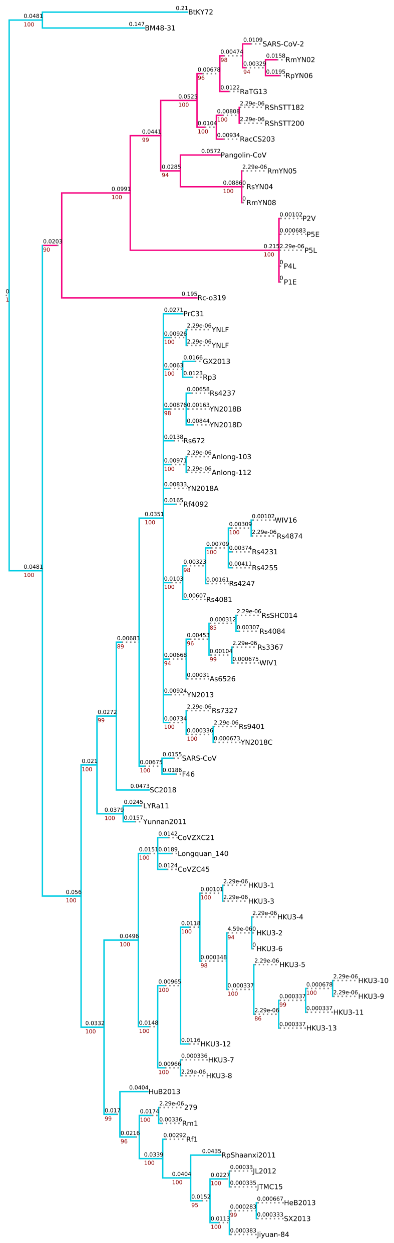


**Figure S1. continued.**

Non-recombinant region 12

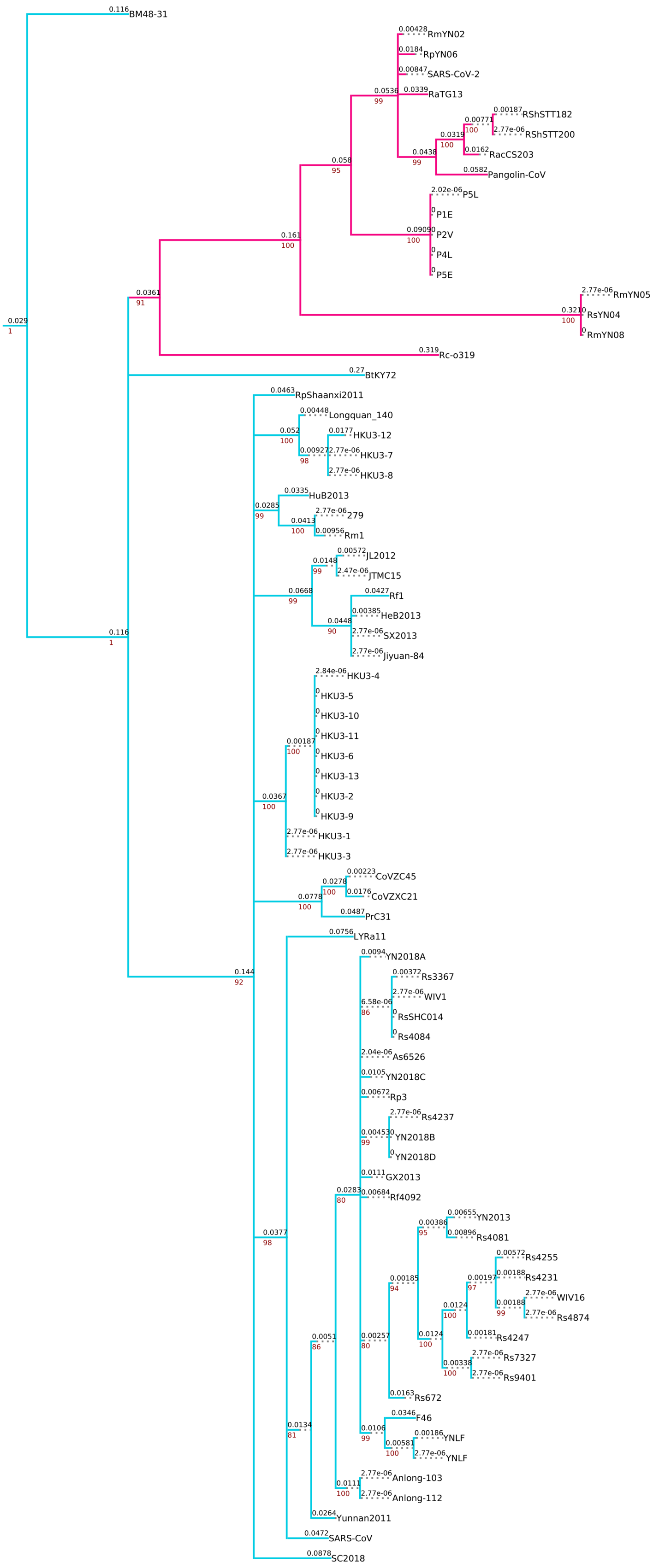


**Figure S1. continued.**

Non-recombinant region 13

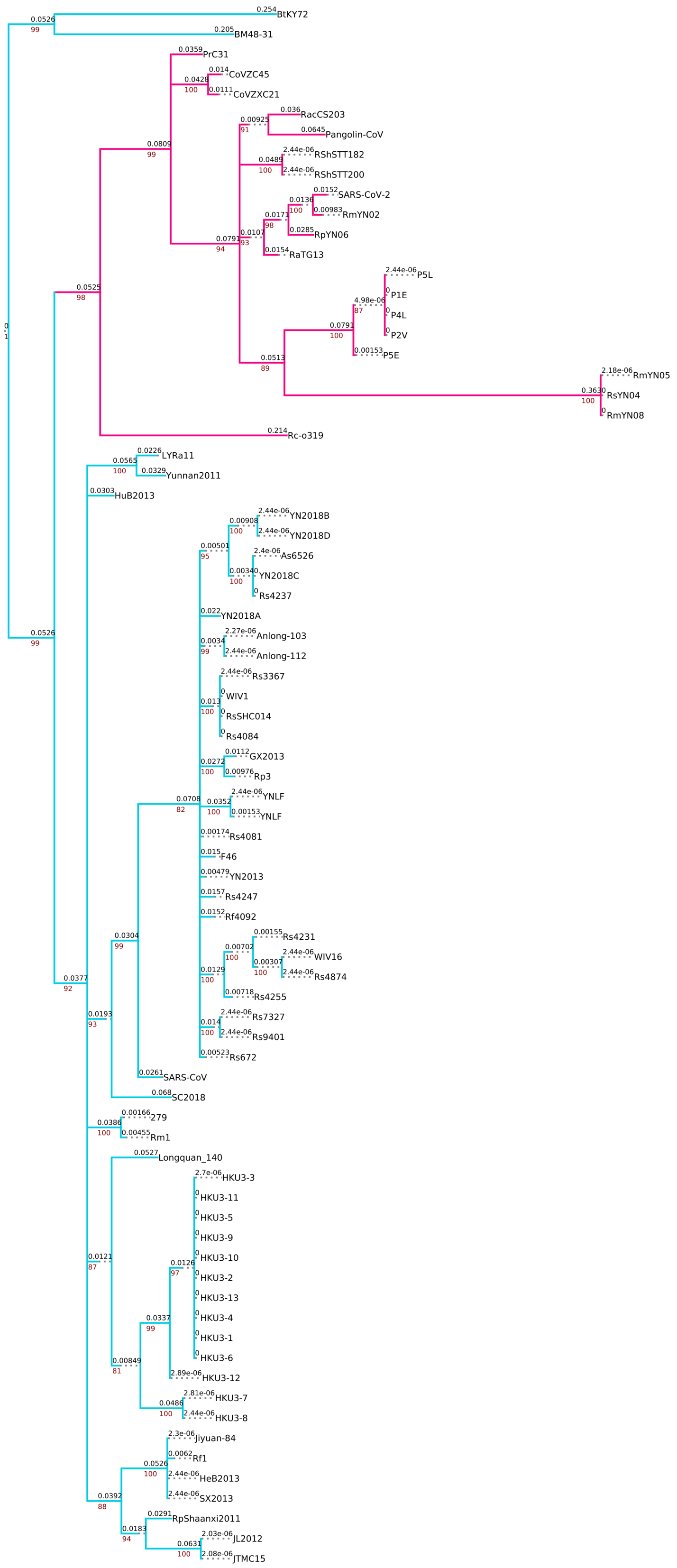


**Figure S1. continued.**

Non-recombinant region 14

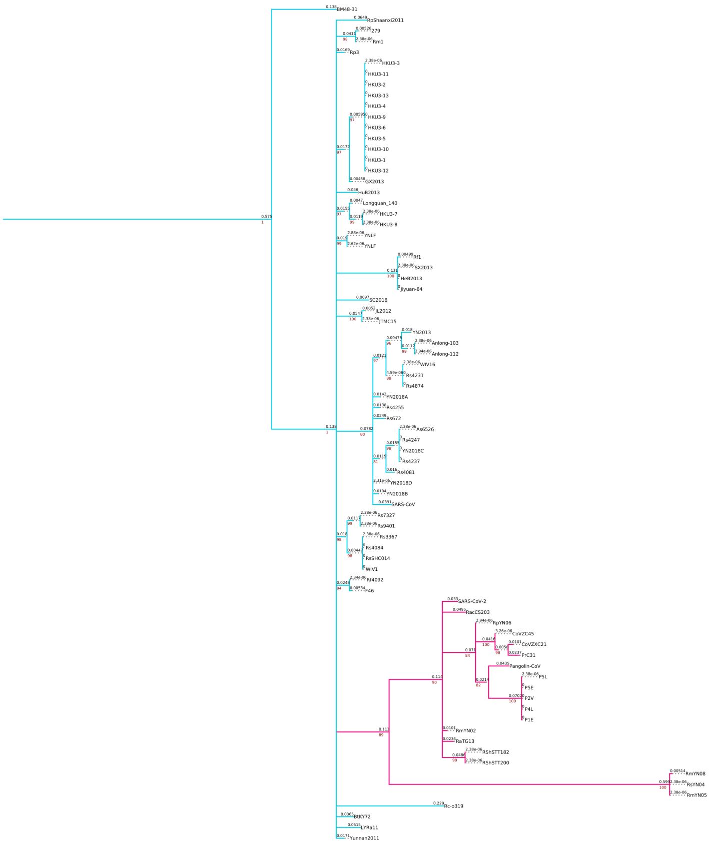


**Figure S1. continued.**

Non-recombinant region 15

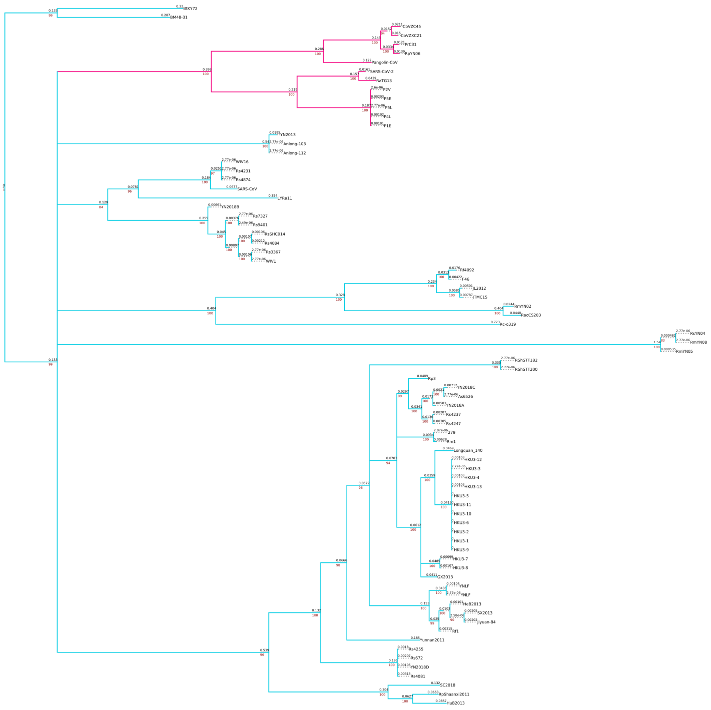

**Figure S1. continued.**

Non-recombinant region 16

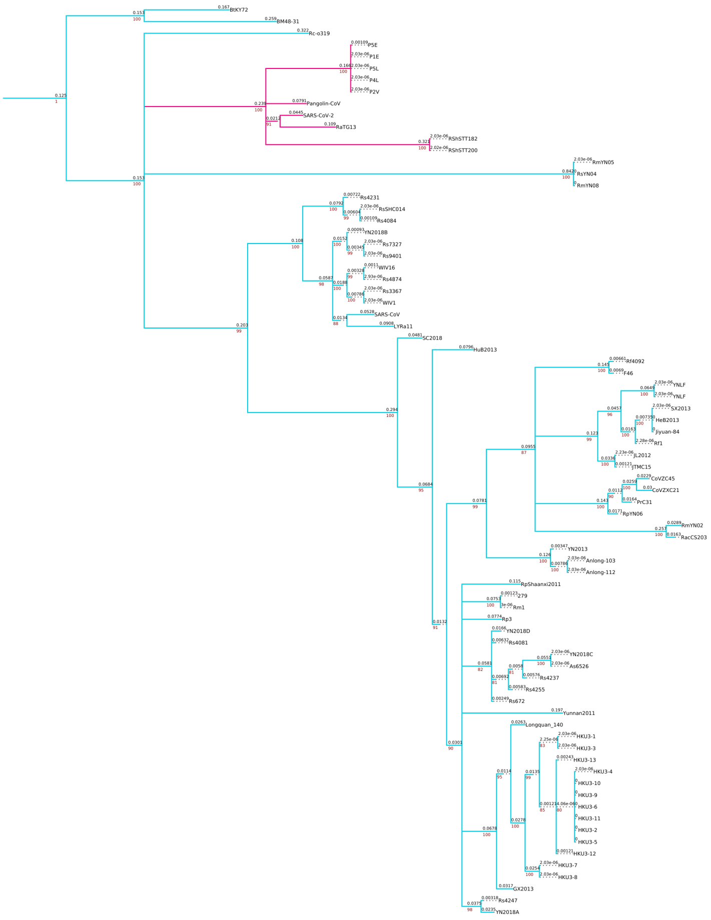

**Figure S1. continued.**

Non-recombinant region 17

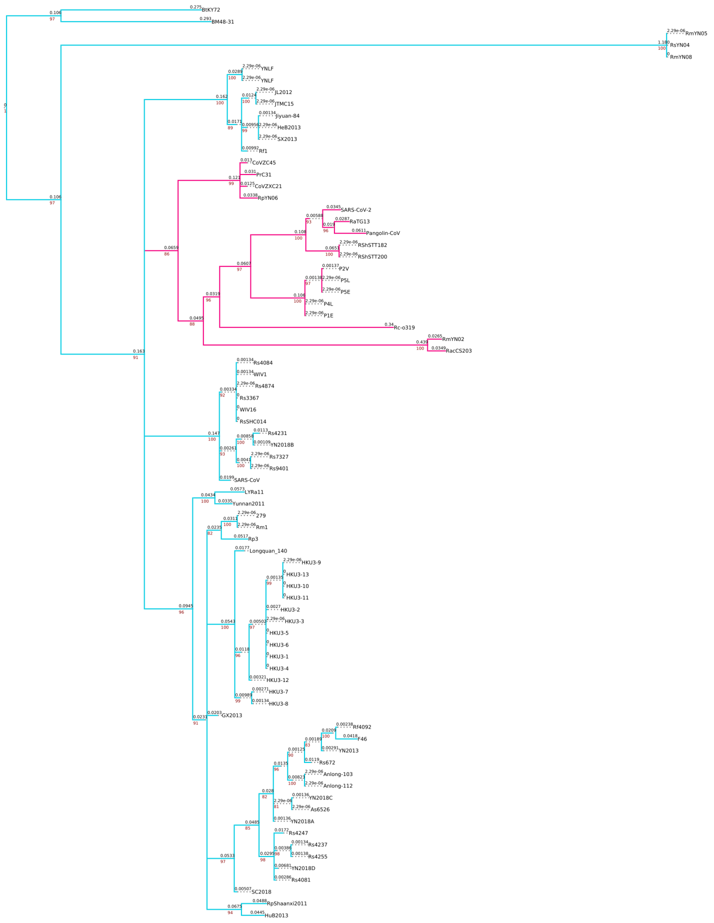


**Figure S1. continued.**

Non-recombinant region 18

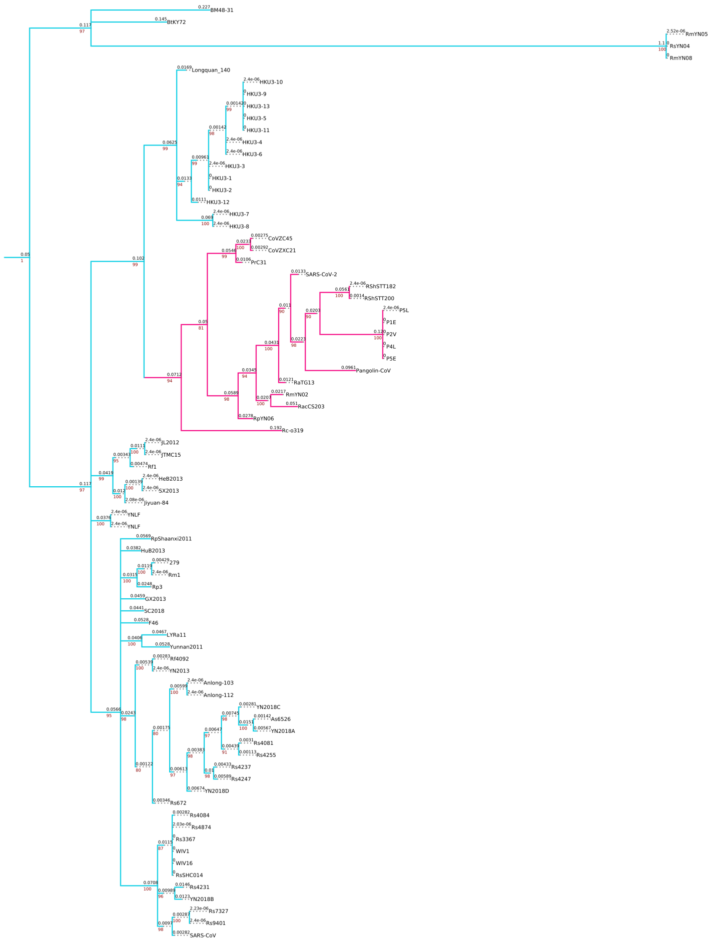

**Figure S1. continued.**

Non-recombinant region 19

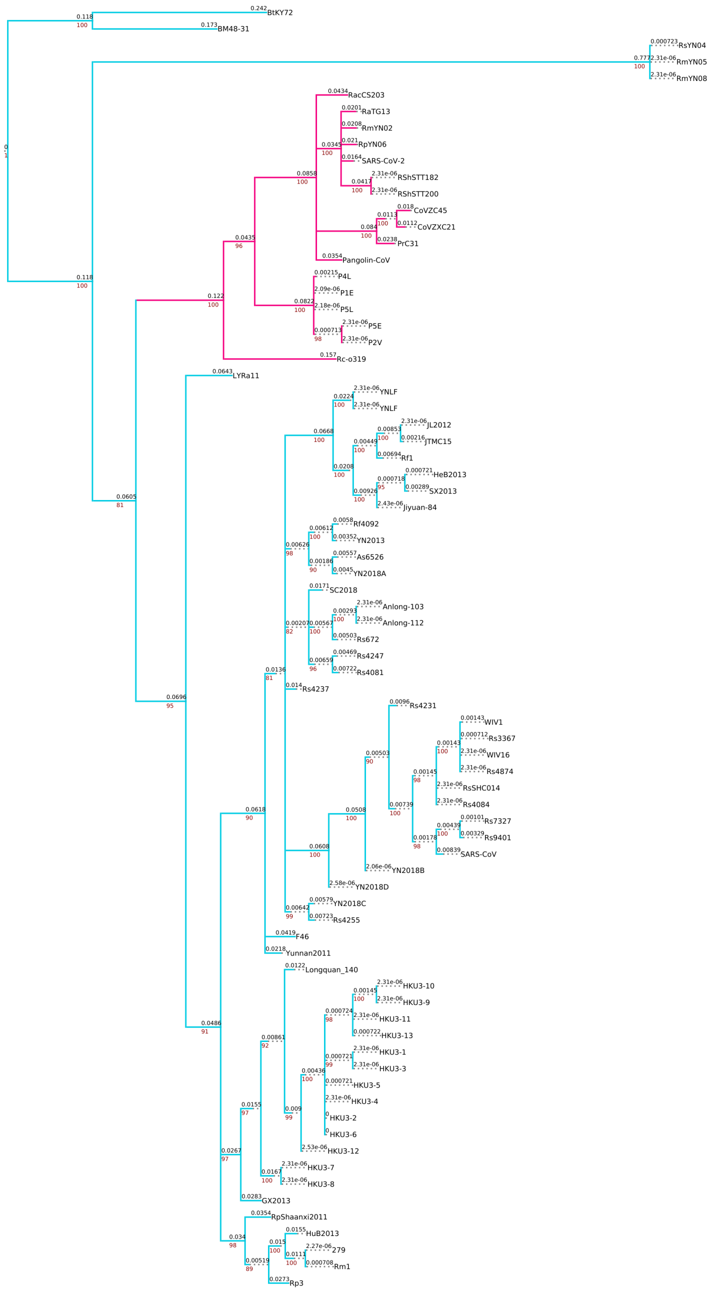


**Figure S1. continued.**

Non-recombinant region 20

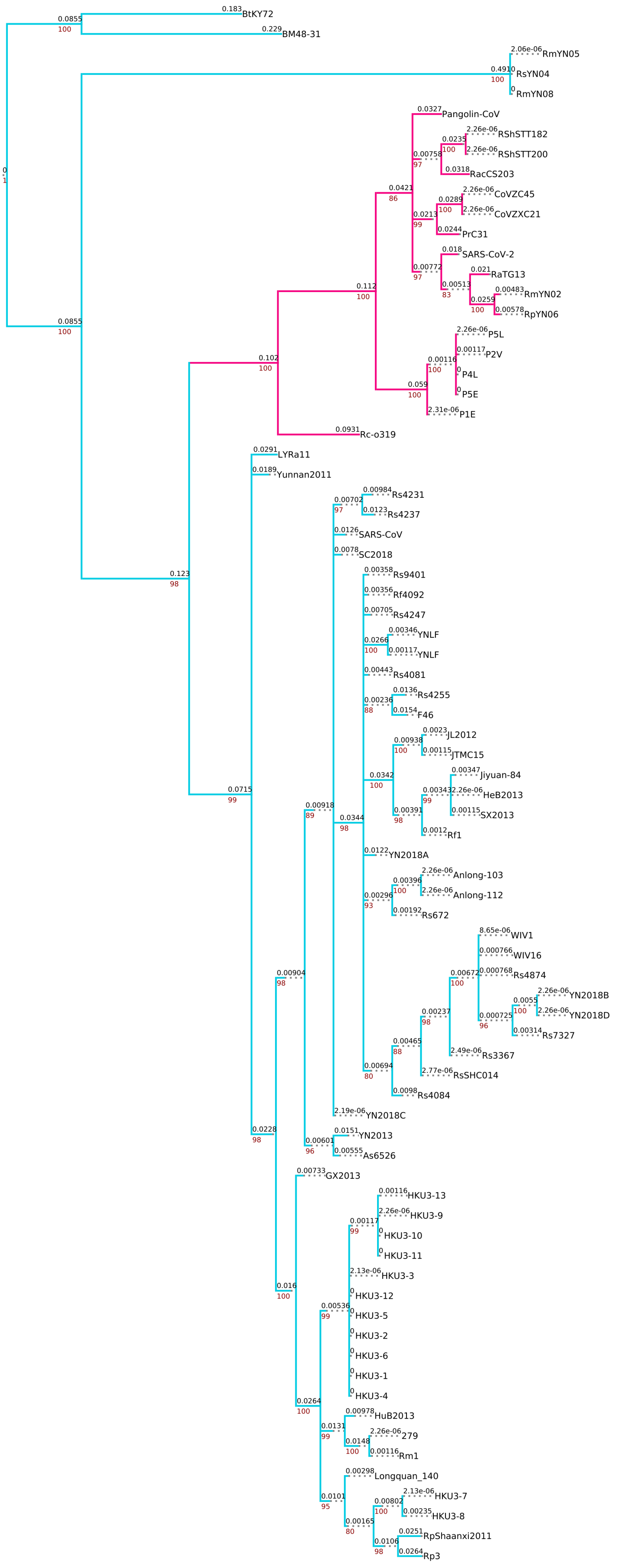


**Figure S1. continued.**

Non-recombinant region 21

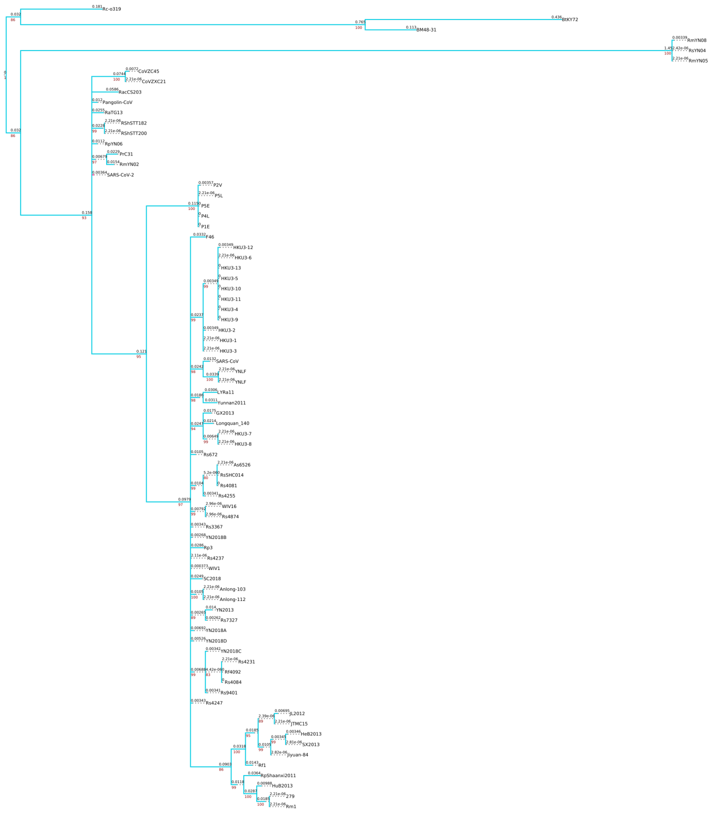

**Figure S1. continued.**

Non-recombinant region 22

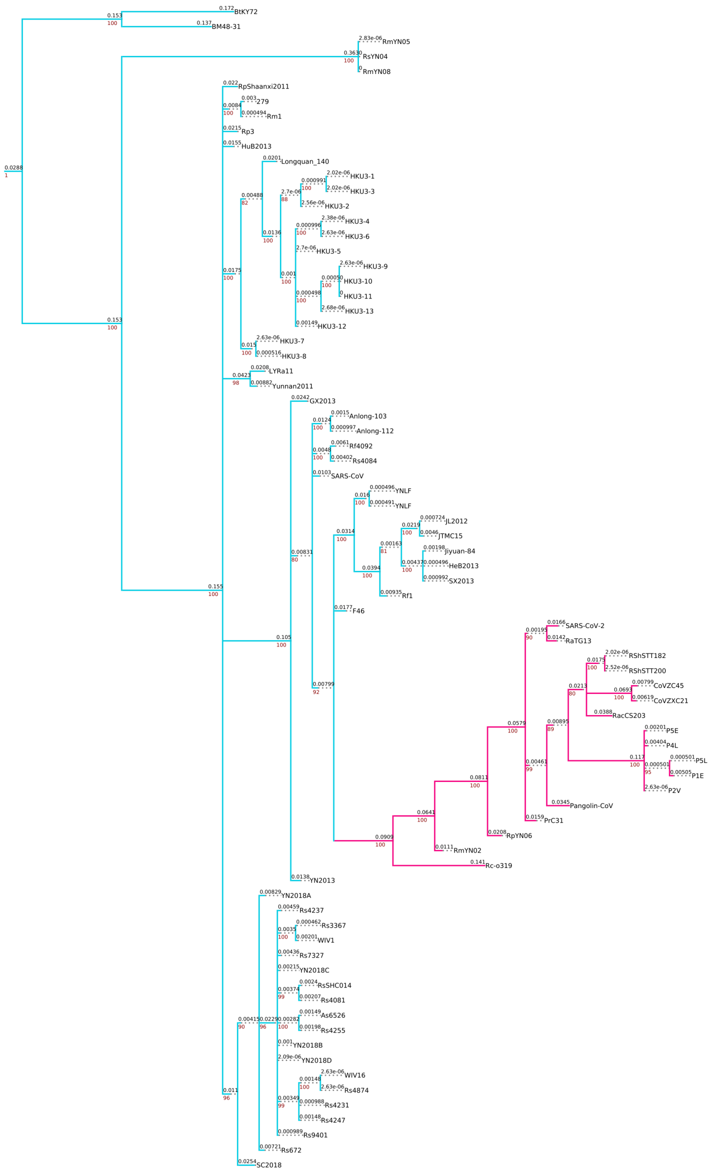


**Figure S1. continued.**

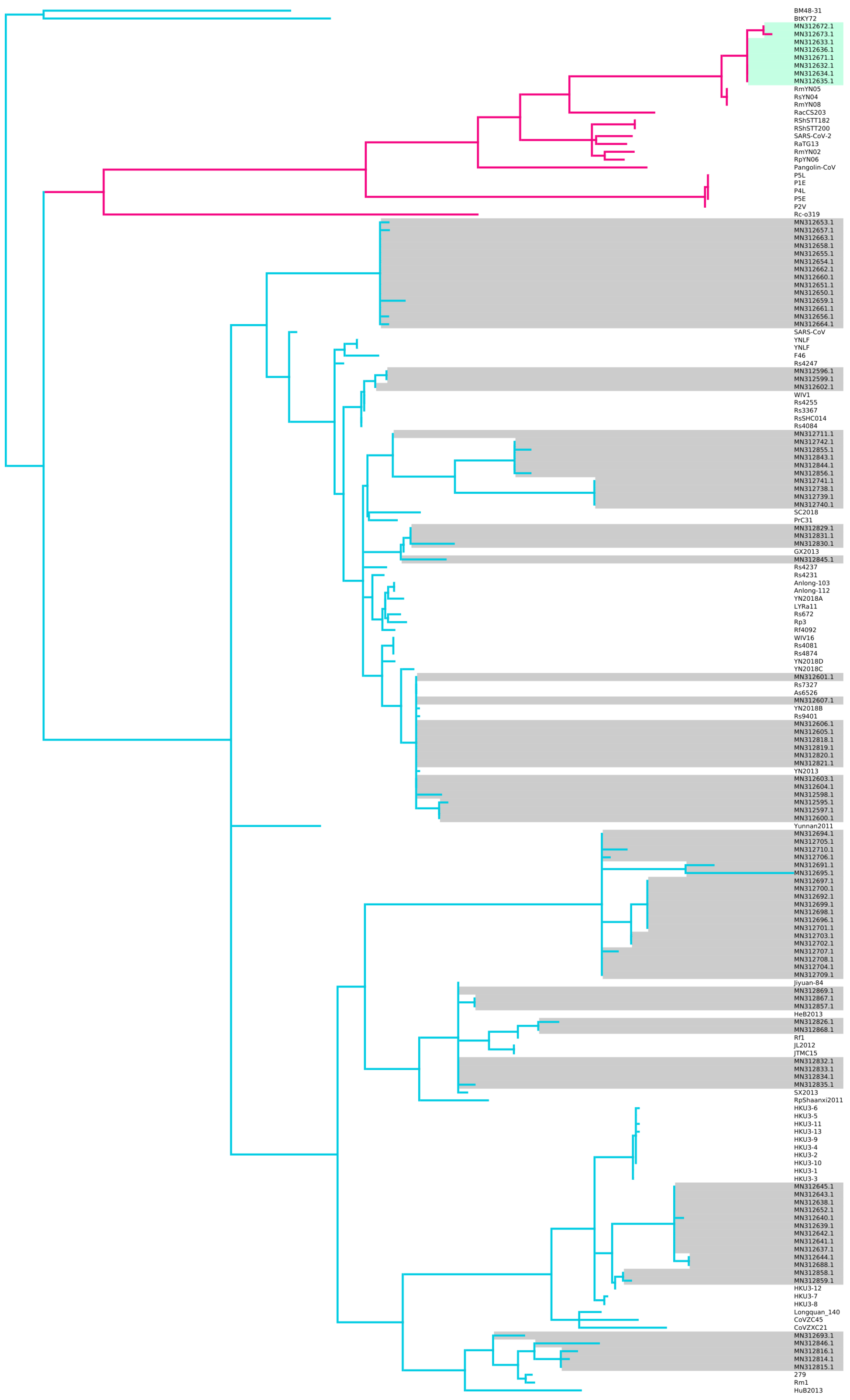


**Figure S2.** Maximum likelihood phylogeny reconstructed using iqtree (GTR+I+4) of all 78 *Sarbecoviruses* used throughout the analysis, including the short RdRp fragments of related *Sarbecoviruses* reported in Latinne *et al.* (2020). The genomic region used for the alignment corresponds to the SARS-CoV-2 reference genome's Wuhan-Hu-1 coordinates 15280 - 16282. Nodes with bootstrap support (10,000 replicates) below 80 have been collapsed. The nCoV clade is annotated in pink and the non-nCoV clade in blue. SARS-CoV-2 and SARS-CoV are highlighted in pink and blue respectively. Viruses from Latinne *et al.* are highlighted in grey, apart from the 7 sequences that cluster within the nCoV clade which are highlighted in green. Out of this cluster of sequences MN312634.1 has been collected from a confirmed R. affinis bat species.
